# Suv420 enrichment at the centromere limits Aurora B localization and function

**DOI:** 10.1101/2021.06.14.448441

**Authors:** Conor P Herlihy, Sabine Hahn, Nicole M Hermance, Elizabeth A Crowley, Amity L Manning

## Abstract

Centromere structure and function are defined by the epigenetic modification of histones at centromeric and pericentromeric chromatin. The constitutive heterochromatin found at pericentromeric regions is highly enriched for H3K9me3 and H4K20me3. While mis-expression of the methyltransferase enzymes, Suv39 and Suv420, that regulate these marks are common in disease, the consequences of such changes are not well understood. Our data show that increased centromere localization of Suv39 and Suv420 suppress centromere transcription and compromise localization of the mitotic kinase Aurora B: decreasing microtubule dynamics and compromising chromosome alignment and segregation. We find that inhibition of Suv420 methyltransferase activity partially restores Aurora B localization to centromeres and that restoration of the Aurora B-containing CPC to the centromere is sufficient to suppress mitotic errors that result when Suv420/H4K20me3 is enriched at centromeres. Consistent with a role for Suv39 and Suv420 in negatively regulating Aurora B, high expression of these enzymes corresponds with increased sensitivity to Aurora kinase inhibition in cancer cells suggesting that increased H3K9 and H4K20 methylation may be an underappreciated source of chromosome missegregation in cancer.

## Introduction

Mitotic chromosome segregation is regulated, in part, through the composition and function of kinetochores, which are protein complexes that link spindle microtubules to chromatin (Thomas et al., 2017, Hinshaw and Harrison, 2018). Kinetochores are, in turn, assembled upon specialized domains of chromatin known as centromeres. Centromeric chromatin is comprised of highly repetitive DNA sequences and is defined epigenetically by the presence of the centromere-specific histone 3 variant CENP-A (Hinshaw and Harrison, 2018, De Rop et al., 2012, Sharma et al., 2019, Ohzeki et al., 2019, Guse et al., 2011, Carroll and Straight, 2006). Flanking the CENP-A containing centromeric chromatin are less ordered repeat sequences that make up the pericentromere. A defining characteristic of pericentromeric chromatin is its enrichment for repressive epigenetic marks, including di- and tri-methylation of lysine 9 on histone H3 and lysine 20 on histone H4 (H3K9me2/3 and H4K20me2/3, respectively), that are strikingly absent from the adjacent centromere (Fioriniello et al., 2020, Sullivan and Karpen, 2004). While CENP-A chromatin is demonstrated to be a critical epigenetic marker upon which the kinetochore is built (De Rop et al., 2012, Barnhart et al., 2011), the function of the epigenetic marks at pericentromeric heterochromatin (PCH) domains remain less clear.

While many studies have implicated changes in heterochromatin in the regulation of genome stability and cancer progression, most focus on the consequences of decreased H3K9me3 and H4K20me3 heterochromatin marks (reviewed in Janssen et al., 2018). Nevertheless, both increased expression of the methyltransferases responsible for placing these repressive marks and decreased expression of demethylases that remove these marks, have been described in various cancer contexts (Janssen et al., 2018, Black et al., 2012, Zhou et al., 2019, Yokoyama et al., 2013). Functional studies in cancer cells indicate that increased levels of H3K9me3 and H4K20me3 contribute to increased motility and metastatic potential (Zhou et al., 2019, Yokoyama et al., 2013) and can limit therapeutic response (Cuellar et al., 2017, Guler et al., 2017).

Studies exploiting a human artificial chromosome (HAC) system have shown that expansion of heterochromatin via direct tethering of Suv39 to the centromere is sufficient to compromise segregation fidelity of the HAC (Ohzeki et al., 2012, Martins et al., 2016). Consistent with these observations, recent work has revealed that expression of Suv39 and Suv420, the methyltransferases responsible for H3K9me2/3 and H4K20me2/3 respectively, correlate with levels of aneuploidy across 10,000+ cancer samples from the Cancer Genome Atlas Project (Taylor et al., 2018). Mitotic segregation errors underlie the generation of aneuploidy and together these data suggest that misregulation of Suv39 and Suv420, and the epigenetic marks they regulate, may be functionally linked to the generation of aneuploidy in cancer.

Key to accurate chromosome segregation is the localization and function of the mitotic kinase Aurora B (Lampson et al., 2004, Liang et al., 2020, Gregan et al., 2011, Krenn and Musacchio, 2015). Aurora B, together with INCENP, Survivin, and Borealin, form the chromosomal passenger complex (CPC). Distributed along chromosome arms at mitotic entry, the CPC re-localizes first to the centromere and kinetochore and later during anaphase to the central spindle (Hindriksen et al., 2017). Concentration of the CPC at the centromere and kinetochore is believed to be crucial for its function in mitotic error correction as it places Aurora B in close proximity to its substrates on kinetochores (Welburn et al., 2010). Aurora B de-stabilizes microtubules by phosphorylating several key regulators of kinetochore-microtubule attachments to promote the removal of merotelic attachments (Cheeseman et al., 2006, Welburn et al., 2010, DeLuca et al., 2011, Cimini et al., 2006, DeLuca et al., 2006). The formation of merotelic attachments during early stages of mitosis are stochastic and loss or functional inactivation of Aurora B kinase activity precludes their correction and corrupts mitotic fidelity (Abe et al., 2016, Broad et al., 2020, Hauf et al., 2003, Huang et al., 2018).

An intricate signaling network involving Haspin-dependent phosphorylation of H3T3 and Bub1-dependent phosphorylation of H2AT120 controls localization of the CPC to the inner centromere. These pathways represent two parallel modes of CPC regulation such that abrogation of either one results in dispersion of the CPC over chromatin and a reduction, but not loss, of CPC at the centromere (Hindriksen et al., 2017, Krenn and Musacchio, 2015, Carretero et al., 2013, Dai et al., 2006, Meppelink et al., 2015, Wang et al., 2010). It remains unclear whether other epigenetic marks at the centromere or pericentromere similarly impact the regulation of CPC localization and function.

## Results

### High expression of Suv39 and Suv420 methyltransferases is prevalent in cancer

Expression data from the TCGA indicate that Suv39 and Suv420 are highly expressed in cancer contexts, with Suv39 isoforms h1 and h2 and Suv420 isoform h2 exhibiting increased average expression compared to corresponding normal tissue in eleven of fourteen cancer subtypes for which paired tumor and normal data is available. (Figure 1A). These enzymes function to regulate constitutive heterochromatin at pericentromeres and in normal cells H3K9me3 and H4K20me3 are enriched near centromeres. In a panel of high Suv420-expressing breast cancer cell lines H4K20me3 is not dispersed uniformly throughout the genome and instead remains enriched near centromeres (Supplemental Figure 1), raising the possibility that Suv420 misexpression may preferentially compromise pericentromere or centromere function and perturb mitotic chromosome segregation. Indeed, recent work describes that high expression of Suv39 or Suv420 positively correlates with pan cancer analyses of increased aneuploidy (Taylor et al., 2018). This relationship is not an artifact of high Suv39 or Suv420 expression in a single cancer type that also happens to be highly aneuploid but instead persists when samples are sorted by cancer subtype. We find that high expression of at least one Suv39 or Suv420 isoform demonstrate a significant positive correlation with aneuploidy in twelve out of twenty cancer contexts represented in the TCGA database (Figure 1B, Supplemental Table 1). Consistent with a correlation with aneuploidy, which is known to correspond with tumor aggressiveness and poor patient outcome (Pfau and Amon, 2012, Weaver et al., 2007, Davoli et al., 2013, Liu et al., 2016), high expression of a Suv39h1/h2 or Suv420h1/h2 isoform corresponds with a significant decrease in disease free survival in four out of twenty cancer subtypes represented in the GEO, EGA and TCGA databases (Supplemental Table 2) (Nagy et al., 2018, Györffy et al., 2010).

**Figure 1.**
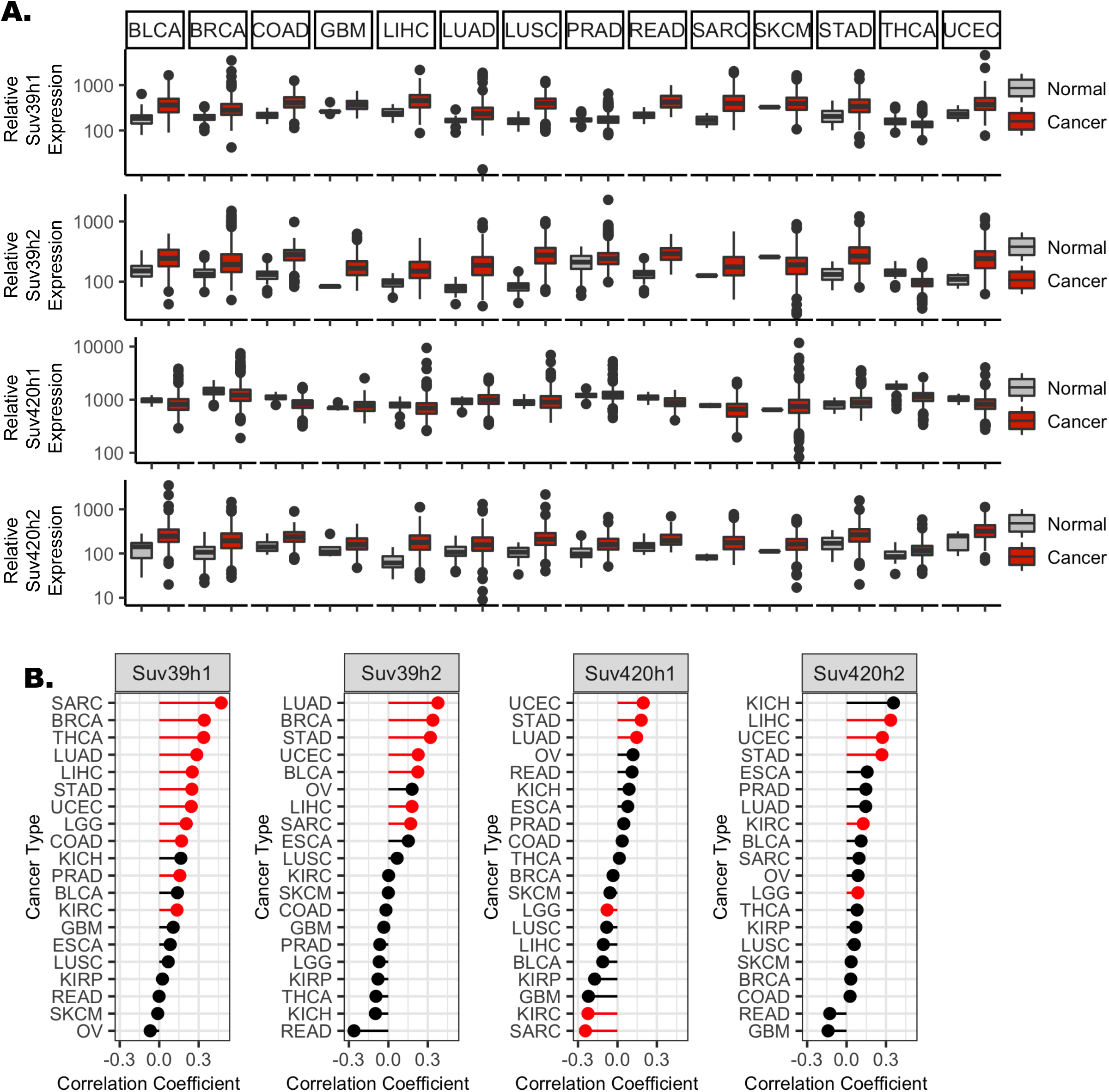
High expression of Suv39 and Suv420 isoforms in cancer correlate with aneuploidy and reduced patient survival. **A)** Gene expression values for Suv39h1, Suv39h2, Suv420h1, and Suv420h2 from TCGA, sorted by cancer subtype indicate increased gene expression compared to normal tissue expression levels in multiple cancer subtypes. **B)** Suv39 and Suv420 isoform expression exhibit a moderate but highly significant correlation with calculated aneuploidy score in several cancer subtypes. Red text indicates significance at p< 0.0025. (See also Supplemental Table 1)

### Depletion of the H3K9me3 demethylase KDM4A compromises mitotic fidelity

To define the consequences of increased H3K9 and H4K20 methylation on mitotic chromosome segregation we used multiple independent approaches to increase the methylated state of H3K9 or H4K20 globally or specifically at the centromere in hTERT-immortalized human retinal pigment epithelial (RPE-1) cells. RPE-1 cells are diploid and genomically stable, exhibiting fewer than one segregation error every 100 cell divisions (Thompson and Compton, 2008). To enhance H3K9 methylation and monitor the impact on mitosis we first utilized siRNA to deplete KDM4A, a demethylase responsible for removing di- and tri-methyl marks from H3K9 (Supplemental Figure 2A) (Berry and Janknecht, 2013). As expected, depletion of KDM4A led to a small but measurable increase in the global level of H3K9 methylation (Supplemental Figure 2D & E). Using time-lapse and fixed cell imaging approaches, we demonstrate that depletion of KDM4A compromises mitotic progression. RPE-1 cells expressing RFP-tagged histone 2B (RFP-H2B) to label chromatin were transfected with non-targeting control (siControl) or KDM4A-targeting siRNA (siKDM4A) for 36 hours and then monitored by live cell fluorescence imaging (Figure 2A). Images were captured every 5 minutes and mitotic cells were tracked to determine the dynamics of chromosome alignment and the timing of anaphase onset. Cells lacking KDM4A exhibit a minor delay in metaphase chromosome alignment and anaphase onset (27.37 +/- 0.40 minutes vs. 34.67 +/- 1.33 minutes, p = 0.007; Figure 2A). Consistent with live cell imaging, immunofluorescence analysis of KDM4A-depleted cells reveal an enrichment in cells where single chromosomes remain near spindle poles when metaphase alignment of the majority of chromosomes has already been achieved (Figure 2B). Although live cell imaging reveals that full metaphase alignment is eventually achieved prior to anaphase onset, nearly 25% of KDM4A-depleted anaphase cells exhibit lagging chromosomes (Figure 2C). Similar results were seen when KDM4A mRNA was targeted for depletion with any one of four independent siRNA sequences (Supplemental Figure 2B, Supplemental Table 3).

**Figure 2.**
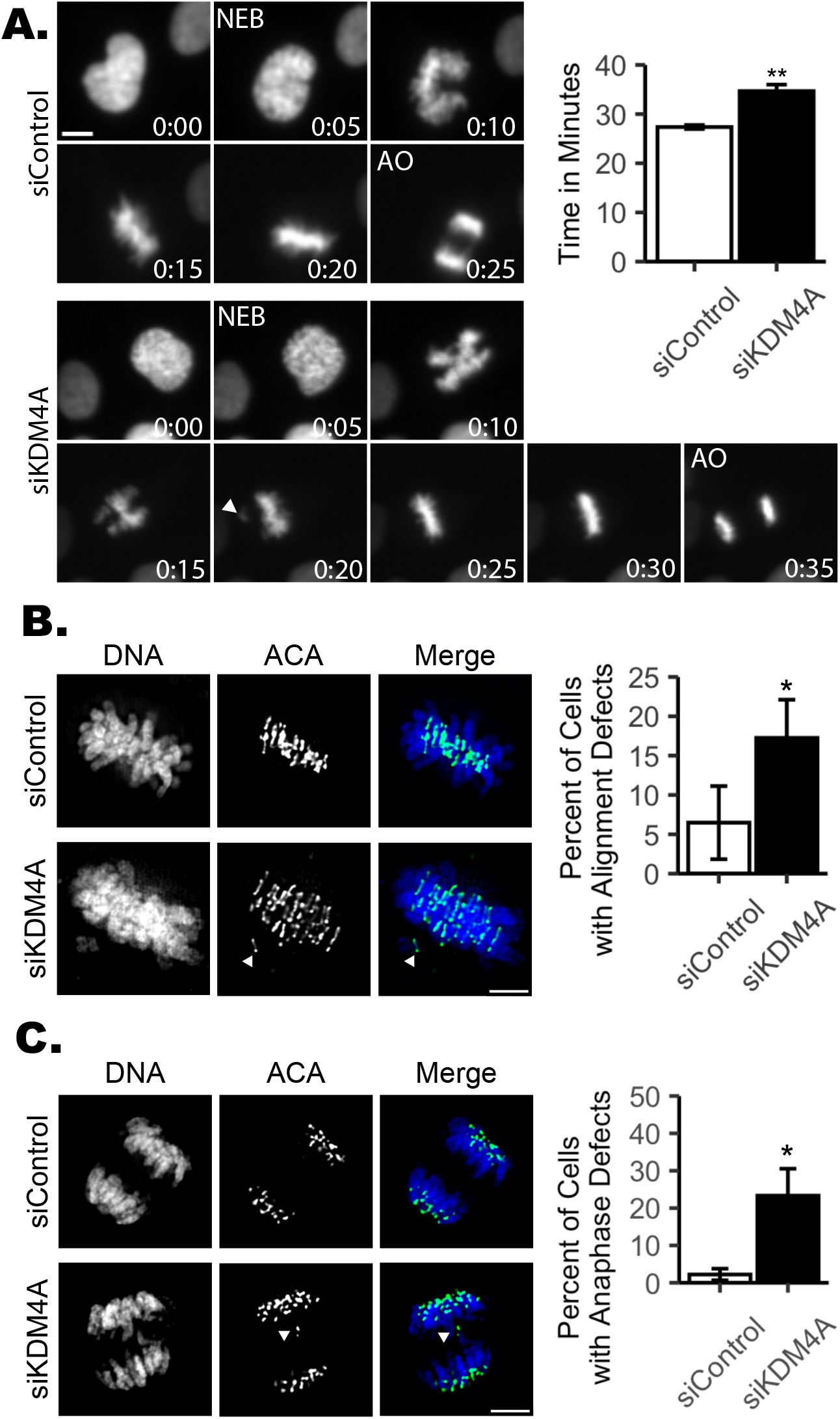
Depletion of KDM4A compromises mitotic fidelity. **A)** Still images and quantification of average mitotic duration from time-lapse microscopy of RFP-H2B expressing RPE-1 cells transfected with either non-targeting control (siControl) or KDM4A-targeting (siKDM4A) siRNAs show cells lacking KDM4A exhibit delayed anaphase onset. Images were captured every 5 minutes. Number insets indicate time progression in minutes. Nuclear envelope breakdown (NEB) and anaphase onset (AO) are indicated. 50 mitotic cells were analyzed per condition for each of 3 biological replicates. **B)** Images and quantification showing KDM4A-depleted mitotic cells have an increase in late prometaphase/metaphase cells that retain 1-3 unaligned chromosomes. **C)** Images and quantification showing KDM4A-depleted anaphase cells exhibit an increase in lagging chromosomes during anaphase. For panels B and C, 30 metaphase or anaphase cells were analyzed per condition for each of 3 biological replicates. For all panels, white arrowheads indicate unaligned or lagging chromosomes; *: p < 0.05, **: p < 0.01. Error bars are +/- SD and statistical analyses were performed between three biological replicates. Scale bars are 5 μm.

### Centromere tethering of the histone methyltransferase Suv39 or Suv420 corrupt mitotic fidelity

As H3K9 methylation is not exclusive to the pericentromere, increased H3K9 methylation that follows KDM4A depletion is not restricted to centromere or pericentromere heterochromatin (Supplemental Figure 2D & E) (Berry and Janknecht, 2013). Therefore, to test if changes in centromere-specific H3K9 methylation levels are sufficient to cause mitotic errors, we engineered RPE-1 cells to inducibly express a GFP-tagged Suv39h1 protein fused to the DNA binding domain of the centromere localized protein CENP-B (cen-Suv39-GFP, Supplemental Figure 2C-E). The DNA binding domain of CENP-B binds to a discrete sequence at the centromere (Pluta et al., 1992), targeting the cen-Suv39-GFP fusion protein to centromeres where it promotes increased H3K9 methylation (Supplemental Figure 2D & E). Comparable to depletion of KDM4A, immunofluorescence imaging of cells following induced expression of cen-Suv39-GFP revealed an increase in metaphase cells with alignment defects and anaphase cells with lagging chromosomes that are not present in the absence of doxycycline-induced expression of the transgene (i.e. Mock vs. Induced: Supplemental Figure 3B, Figure 3A).

**Figure 3.**
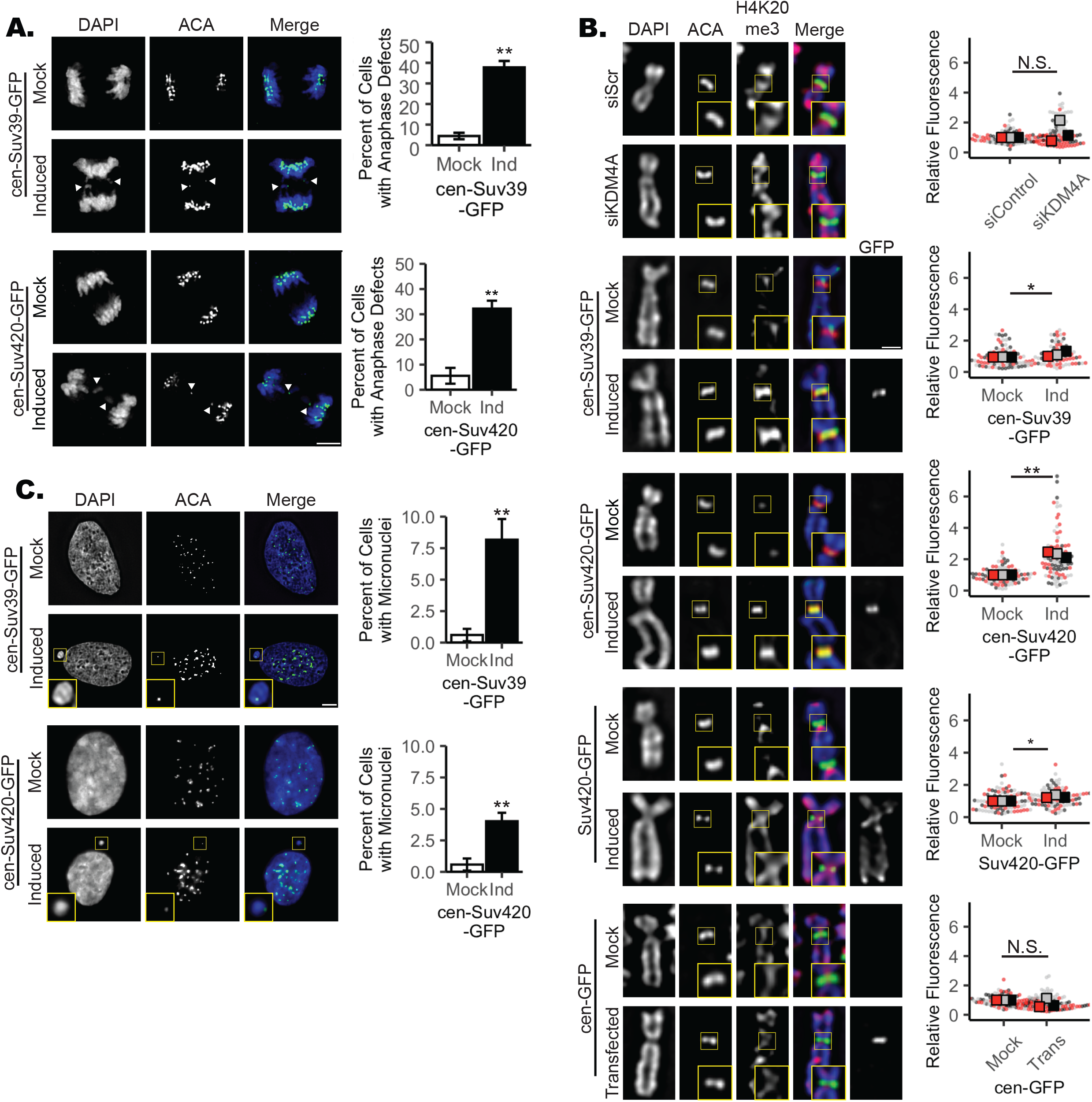
Increased H4K20me3 corresponds with decreased mitotic fidelity. **A)** Images and quantification showing an increase in lagging chromosomes during anaphase following induced expression of cen-Suv39-GFP or cen-Suv420-GFP. White arrowhead indicates a lagging chromosome. Scale bar is 5 μm. **B)** Images and quantification showing induction of cen-Suv420-GFP promotes a substantial increase in centromere-localized H4K20me3 levels, while siKDM4A, cen-Suv39-GFP, and (nontethered) Suv420-GFP induce only a moderate changes in H4K20me3 at centromeres. Cen-GFP does not alter centromere H4K20me3 levels. Insets are 1.5x enlargements of ACA-stained kinetochore pairs. Scale bar is 1 μm. **C)** Images and quantification showing an increase in micronuclei formation following induction of cen-Suv39-GFP or cen-Suv420-GFP. Insets are 3X enlargements of individual micronuclei. Presence of an ACA-labeled kinetochore within each micronucleus indicates they likely form following mitotic segregation errors. Scale bar is 5 μm. Alignment and segregation errors were assessed in 30 cells per condition, per replicate. Intensity analyses were done on 10 kinetochores for each of 30 cells per condition, per replicate. For micronuclei analyses, a minimum of 240 cells were scored per condition, per replicate. All statistical analyses were performed between three biological replicates, *: p < 0.05, **: p < 0.01. Error bars are +/- SD.

H3K9 methylation promotes recruitment and binding of heterochromatin protein 1 (HP1). HP1 in turn serves as a platform to recruit a number of factors to heterochromatin, including the H4K20 methyltransferase Suv420 (Janssen et al., 2018, Hahn et al., 2013, Schotta et al., 2004). Thus, establishment of H3K9me3 by Suv39 is indirectly implicated in deposition of H4K20me3. Consistent with this relationship, we find that centromere targeting of Suv39 (the H3K9me2/3 methyltransferase) enhances H4K20me3 levels at the centromere (Figure 3B). To delineate the roles of H3K9 and H4K20 centromere methylation in the regulation of mitotic progression we next expressed a GFP-tagged Suv420h2 protein fused to the DNA binding domain of centromere protein CENP-B (cen-Suv420-GFP). Cen-Suv420-GFP localizes to centromeres and is sufficient to promote a marked enrichment of H4K20me3 at centromeres without altering H4K20me3 levels along chromosome arms (Figure. 3B, Supplemental Figure 3C). As seen with KDM4A depletion and cen-Suv39-GFP expression, induced expression of cen-Suv420-GFP leads to metaphase alignment defects (Supplemental Figure 3B) and anaphase segregation errors (Figure 3A), suggesting that increased H4K20me3 at centromeres is sufficient to corrupt kinetochore regulation. Chromosomes that lag during anaphase can be excluded from the main nucleus when the nuclear envelope reforms, resulting in the formation of a separate micronucleus. Consistent with the presence of chromosome segregation defects, cells expressing cen-Suv39-GFP or cen-Suv420-GFP also exhibit an increase in the number of interphase cells within the population that have centromere-positive micronuclei (Figure 3C).

### Centromere enrichment of Suv420 compromises mitotic error correction mechanisms

Formation of syntelic and merotelic microtubule attachments, where both kinetochores of a chromosome pair are bound by microtubules from the same spindle pole, occur stochastically during mitosis. Error correction mechanisms to destabilize such mal-attachments persist throughout prometaphase and metaphase (Cimini et al., 2003). Delays in chromosome alignment to the metaphase plate, and the presence of lagging chromosomes during anaphase, can result from persistent merotelic attachments and may indicate that the number of merotelic attachments formed in mitosis overwhelm the error correction machinery (Cimini et al., 2003). Imbalances in error corrective mechanisms that result in merotelic attachments and segregation errors arise due to either an increased burden in the number of merotelic attachments formed, or from an underlying defect in the error correction machinery. To determine if cells with altered centromere methylation are predisposed to forming merotelic attachments or instead are deficient in correcting these mal-attachments, KDM4A-depleted cells or those induced to express either cen-Suv39-GFP or cen-Suv420-GFP were exposed to the Eg5 inhibitor monastrol for 4 hours (Figure 4A). Inhibition of the mitotic kinesin Eg5 causes spindle poles to collapse, resulting in a monopolar spindle structure. These spindles cannot form amphitelic chromosome attachments and instead form syntelic or monotelic attachments. Upon washout of monastrol, a large portion of kinetochore attachments are converted to merotelic attachments (Kapoor et al., 2000, Lampson et al., 2004). Destabilization of mal-attached microtubules is a prerequisite for acquisition of proper kinetochore biorientation and chromosome alignment such that the time needed to achieve complete metaphase alignment following monastrol washout is an indication of error correction efficiency. Immunofluorescence imaging demonstrates that, following monastrol washout, bipolar spindles form and the majority of control cells restore chromosome biorientation and achieve metaphase alignment within 40 minutes. In contrast, though spindle bipolarity is achieved similarly to control cells, KDM4A-depleted, cen-Suv39-GFP expressing and cen-Suv420-GFP expressing cells all exhibit a delayed progression to metaphase, with fewer than half of all cells able to achieve metaphase alignment within the same time frame as control cells (Figure 4A). Together, these data indicate that the mitotic error correction machinery is compromised under conditions where H3K9me3 or H4K20me3 is increased.

**Figure 4.**
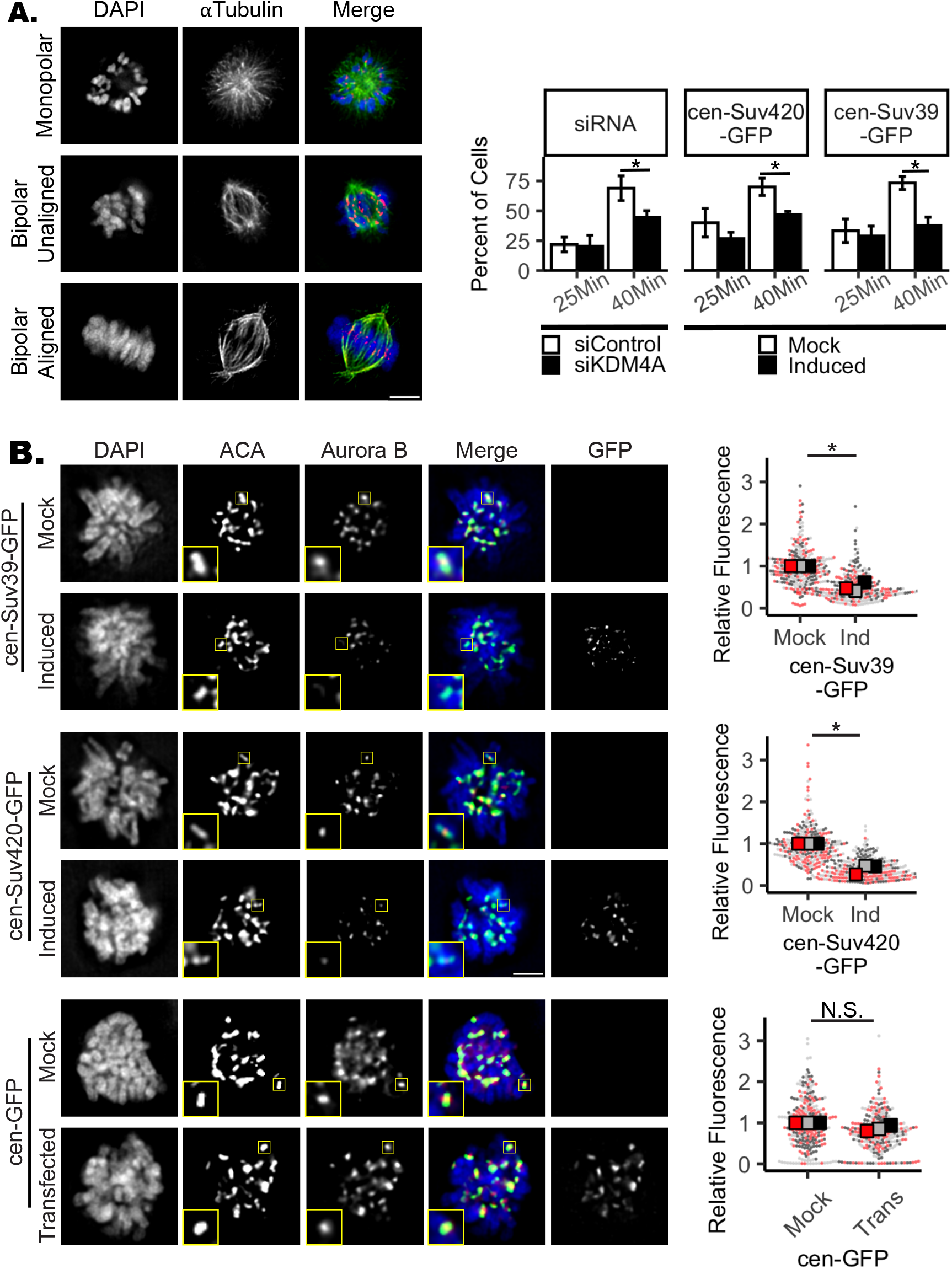
Mitotic error correction and Aurora B localization are compromised when centromere Suv420/H4K20 methylation levels are increased. **A)** Images and quantification showing progression to full metaphase alignment following inhibition of Eg5 with monastrol and subsequent drug washout. Cells lacking KDM4A or expressing cen-Suv39-GFP or cen-Suv420-GFP are delayed in achieving metaphase alignment, indicating a deficiency in mitotic error correction capacity. **B)** Images and quantification of Aurora B kinase localization at centromeres of nocodazole-treated prometaphase cells. Induced expression of cen-Suv39-GFP or cen-Suv420-GFP, but not cen-GFP, reduced Aurora B intensity at centromeres. Insets are 2X enlargements of single ACA-stained kinetochore pairs. Scale bars are 5 μm. Kinetochore analyses were done on 10 kinetochores for each of 30 cells per condition, per replicate. Alignment was assessed in 50 cells per condition, per replicate. For all panels statistical analyses were performed between three biological replicates, *: p < 0.05, **: p < 0.01. Error bars are +/- SD.

### Centromere localization of the CPC is sensitive to centromere enrichment of Suv39 or Suv420

Aurora B is a mitotic kinase that functions as a master regulator of error correction. Loss or functional inactivation of Aurora B kinase activity leads to persistent merotelic attachments and segregation errors (Lampson et al., 2004, Liang et al., 2020, Gregan et al., 2011, Krenn and Musacchio, 2015, Kallio et al., 2002, Hauf et al., 2003, Ditchfield et al., 2003). To determine if changes in Aurora B, localization and/or function underlie observed defects in mitosis following manipulations that increase H3K9me3 and/or H4K20me3, we first performed quantitative immunofluorescence to measure Aurora B and INCENP (a component of the Aurora B-containing CPC complex) recruitment to centromeres following expression of either cen-Suv39-GFP or cen-Suv420-GFP, or depletion of KDM4A. By measuring pixel intensity of Aurora B or INCENP antibody staining across kinetochore pairs stained with anti-centromere antigen (ACA) in metaphase and/or nocodazole-arrested prometaphase cells we identified and assessed Aurora B localization at centromeres and kinetochores. We find that the intensity of both Aurora B and INCENP at mitotic centromeres is significantly reduced following depletion of KDM4A or induced expression of either cen-Suv39-GFP or cen-Suv420-GFP (Figure 4, Supplemental Figure 4). Aurora B, Borealin, INCENP, and Survivin are all expressed at similar levels in cells with and without induction of cen-Suv39-GFP or cen-Suv420-GFP, suggesting that decreased staining intensity is not a consequence of changes in CPC expression (Supplemental Figure 4C). Importantly, expression of centromere targeted GFP alone (cen-GFP) or non-targeted Suv420 (Suv420-GFP), conditions that do not enhance centromere enrichment of H4K20me3, do not perturb centromere levels of Aurora B (Figure 4, Supplemental Figures 3 and 4). Instead, reduction in centromere localization of CPC components is sensitive to Suv420 and/or H4K20me3 at centromeres as treatment with A196, a specific small molecule inhibitor of Suv420 methyltransferase activity (Bromberg et al., 2017), is sufficient to partially restore Aurora B localization to centromeres and to promote efficient chromosome alignment along the metaphase plate (Figure 5).

**Figure 5.**
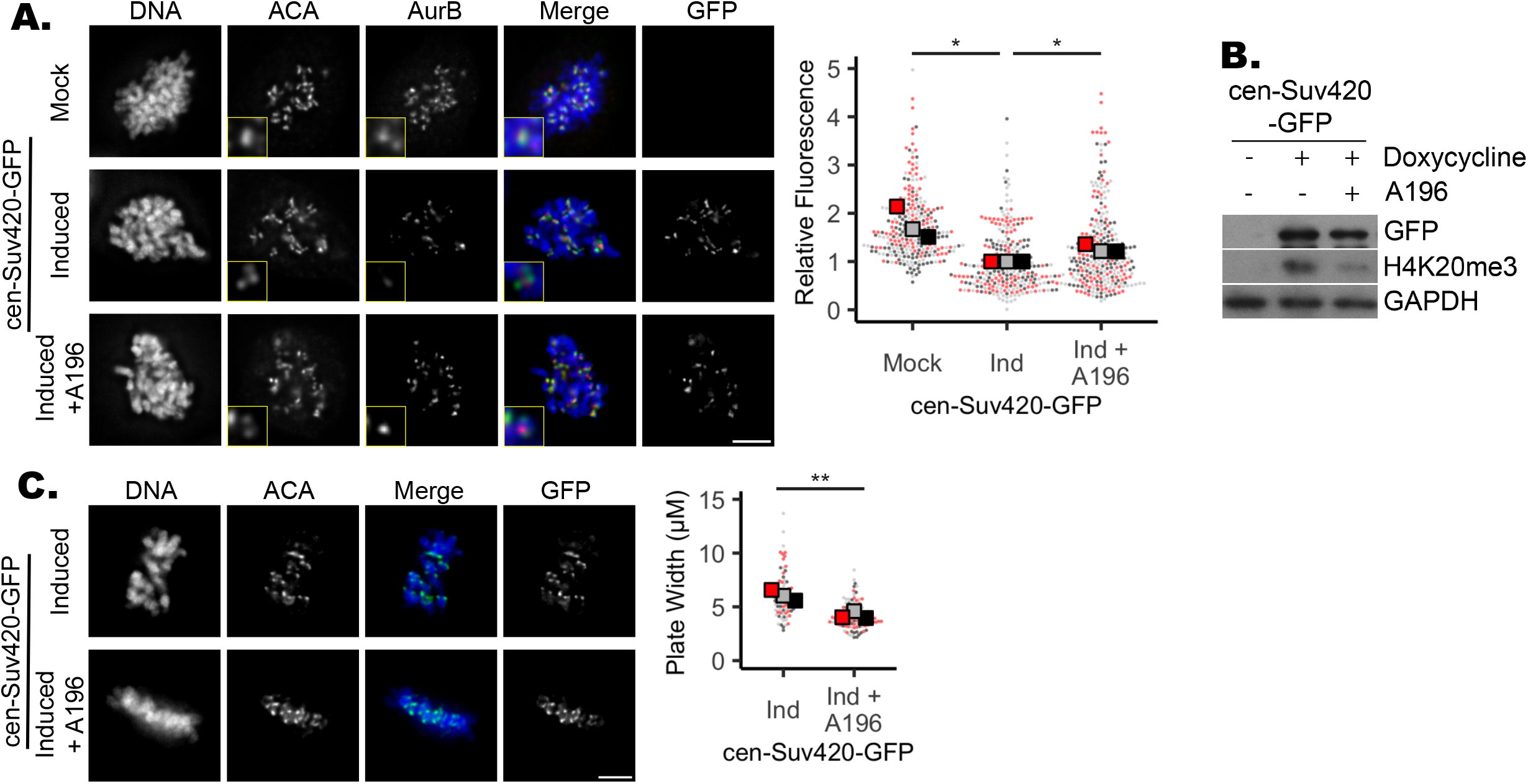
Inhibition of Suv420 methyltransferase activity restores Aurora B localization and corrects metaphase alignment defects. **A & B)** Treatment of cen-Suv420-GFP expressing cells with 200 nM Suv420 inhibitor A196 reduces H4K20me3 levels and partially restored Aurora B localization to the centromere. **C)** Chromosome alignment defects seen in cells expressing cen-Suv420-GFP are suppressed following concurrent treatment with A196. Insets are 2X enlargements of single ACA-stained kinetochore pairs. Kinetochore analyses were done on 3 kinetochores for each of 30 cells per condition, per replicate. Alignment was assessed in 30 cells per condition, per replicate. For all panels statistical analyses were performed between three biological replicates, *: p < 0.05, **: p < 0.01.

Phosphorylation of Hec1 (at serine 55) by Aurora B de-stabilizes kinetochore microtubule attachments to permit error correction. Consistent with compromised Aurora B localization, we find centromeres in mitotic cells have decreased Hec1 phosphorylation, but not overall levels of Hec1, when cen-Suv39-GFP or cen-Suv420-GFP is expressed (Figure 6A & B). These cells also exhibit a corresponding increase in stable microtubules that are resistant to cold-induced depolymerization (Figure 6C). A similar decrease is seen in phosphorylation levels of Aurora B target CENP-A (at serine 7) at centromeres (Supplemental Figure 5A & B). However, consistent with other conditions that specifically perturb Aurora B activity at centromere but not along chromosome arms (Wang et al., 2010), no change is seen in the phosphorylation of H3 (at serine 10), a substrate of Aurora B on chromosome arms (Supplemental Figure 5C).

**Figure 6.**
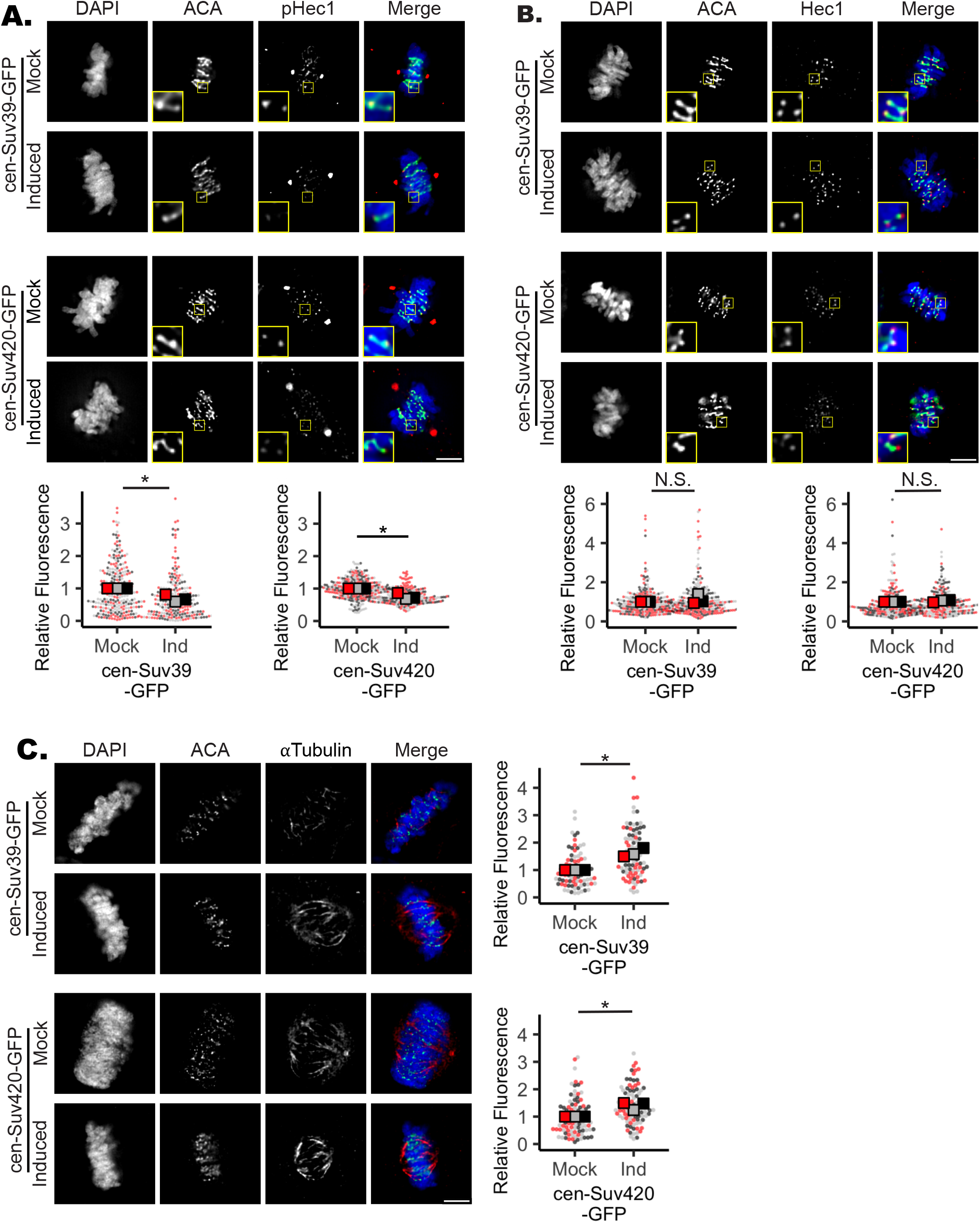
Increased H4K20me3 reduces phosphorylation of Aurora B target Hec1 and increases microtubule stability. **A & B)** Images and quantification showing that pHec1, but not total Hec1 levels at the centromere are decreased following cen-Suv39-GFP or cen-Suv420-GFP expression. Note that this pHec1 antibody stains both centrosomes and kinetochores. Insets are 3X enlargements of single ACA-stained kinetochore pairs. Staining intensity between ACA peaks was measured for 3 kinetochore pairs in each of 30 metaphase cells per condition, in each of 3 biological replicates. **C)** Images and quantification showing that induction of cen-Suv39-GFP or cen-Suv420-GFP expression increases the resistance of kinetochore microtubules to cold-induced depolymerization, indicating an increase in stability of kinetochore-microtubule attachments. Microtubule intensity was measured in each of 30 metaphase cells per condition, in each of 3 biological replicates. *: p < 0.05, **: p < 0.01. Scale bars are 5 μm.

Aurora B localization to centromeres is influenced by several histone modifications, including phosphorylation of threonine 120 on histone 2A (H2A-T120p) by BUB1 and phosphorylation of threonine 3 on histone 3 (H3-T3p) by Haspin. Each of these histone phosphorylation events is independently sufficient to recruit Aurora B to centromeres (Liang et al., 2020, Broad et al., 2020, Hadders et al., 2020). However, we do not see reduction in either of these histone marks in contexts where H4K20me3 has been directly or indirectly increased (Supplemental Figure 6A & B), suggesting indirect changes in these known regulatory marks cannot explain decreased centromere CPC complex localization.

Localization of the CPC is also sensitive to centromere transcription. Centromere transcripts have been shown to bind the CPC and regulate both CPC centromere localization and Aurora B activation (Jambhekar et al., 2014, Blower, 2016, Ideue et al., 2014). To test whether centromere transcription is altered by centromere tethering of Suv39-GFP or Suv420-GFP we measured levels of centromere transcripts in nocodazole-synchronized cells following 24 hours of cen-Suv39-GFP or cen-Suv420-GFP expression. While qPCR analysis shows expression of centromeric αsatellite RNA is readily detected during mitosis, levels of each transcript were reduced by roughly half in mitotic cells expressing cen-Suv39-GFP or cen-Suv420-GFP (Supplemental Figure 6C). Consistent with the centromere tethering of these constructs, suppression of transcription appears restricted to centromeres as transcripts from non-centromere regions, such as housekeeping genes (GAPDH and actin) and CPC components remain unchanged in these samples (Supplemental Figure 4C). These data suggest that suppression of transcription underlies defects in Aurora B localization when centromere levels of Suv39/H3K9me3 and/or Suv420/H4K20me3 are increased.

### Expression level of Suv39 and Suv420 correspond with sensitivity to Aurora kinase inhibition in cancer cell lines

Decreased CPC localization and increased rates of chromosome segregation errors may render cells exquisitely sensitive to further inhibition of Aurora kinase activity. To test this possibility, cells with and without induction of cen-Suv39-GFP or cen-Suv420-GFP expression were treated with inhibitors targeting Aurora B Kinase (Barasertib), the related Aurora A kinase (Alisertib), or the mitotic kinase MPS1 and monitored for mitotic fidelity. While control, cen-Suv39-GFP and cen-Suv420-GFP expressing cells were all similarly sensitive to MPS1 inhibition, we find that cells expressing cen-Suv39-GFP or cen-Suv420-GFP are susceptible to increased anaphase lagging chromosomes following short term treatment with low nanomolar concentrations of both Aurora kinase inhibitors, while control cells are not (Figure 7A). We next tested whether mitotic defects in cen-Suv420-GFP expressing cells could be suppressed by concurrently tethering the Aurora B containing CPC to the centromere. Cen-Suv420-GFP expressing cells were monitored for anaphase defects following expression of cen-INCENP-mCherry (Wang et al., 2011). Cells expressing cen-Suv420 exhibit a high rate of lagging chromosomes during anaphase (Figures 3A & 7E). The frequency of these defects was reduced when the cen-INCENP-mCherry construct was expressed simultaneously. Importantly, the level of cen-Suv420-GFP at centromeres was not significantly altered following cen-INCENP-mCherry expression (Figure 7B-D), suggesting this rescue is not the result of a competition between INCENP and Suv420 for centromere binding.

**Figure 7.**
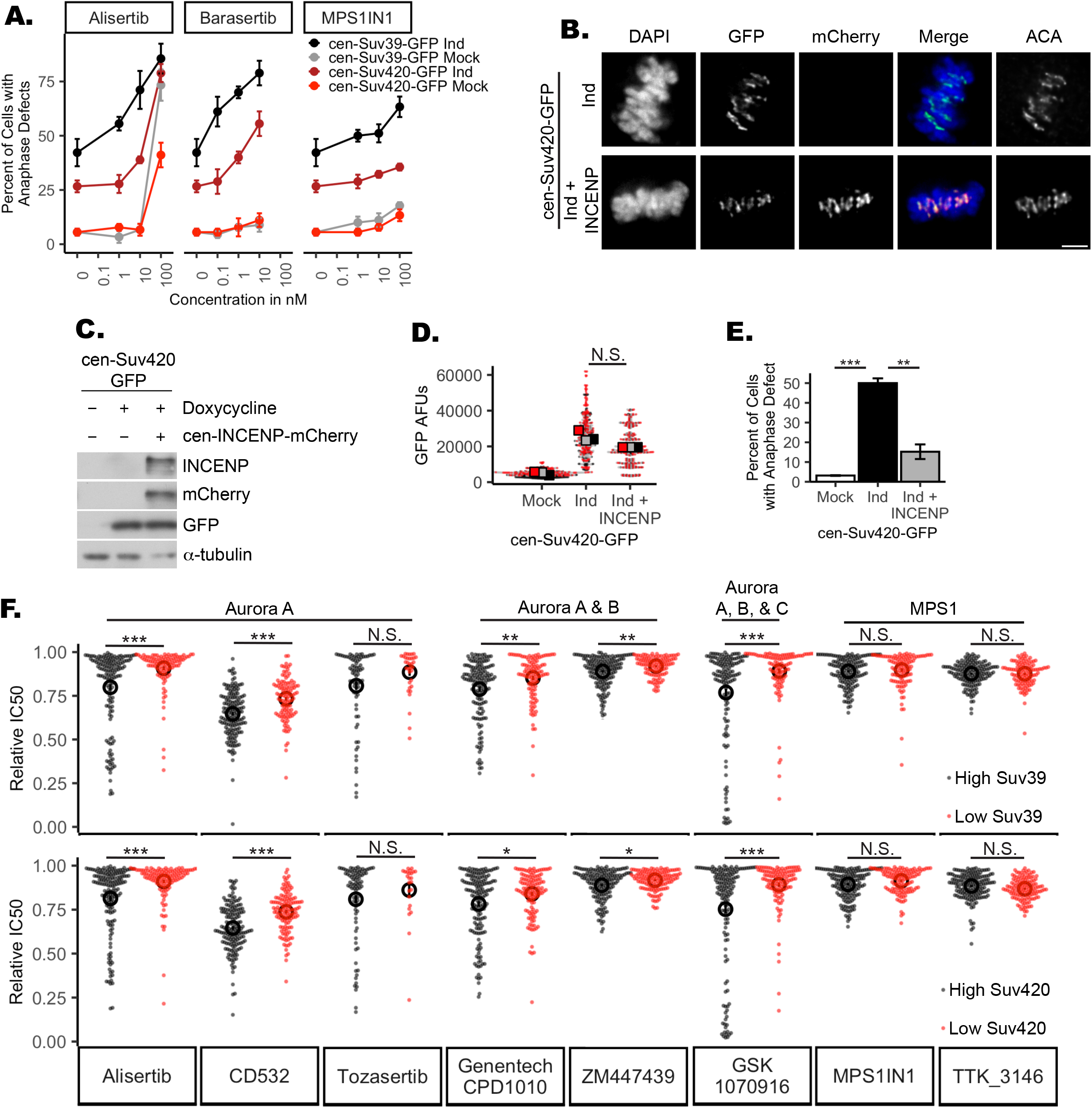
Expression of Suv39 and Suv420 sensitizes cells to Aurora kinase inhibition. **A)** Quantification of anaphase defects showing that cells induced to express cen-Suv39 or cen-Suv420 exhibit an increase in anaphase lagging chromosomes following low nanomolar concentrations of Aurora kinase inhibition while their uninduced counterparts do not. Cells both with and without induction of cen-Suv39-GFP or cen-Suv420-GFP are similarly sensitive to inhibition of the mitotic kinase MPS1. (+/- cen-Suv39-GFP induction: p= 0.00003, 0.0003, and 0.844 for Alisertib, Barasertib, and MPS1IN1, respectively; +/-cen-Suv420-GFP: p = 0.003, 0.004, and 0.724 for Alisertib, Barasertib, and MPS1IN1, respectively) Error bars are +/- SD. **B & C)** Cen-INCENP-mCherry localizes to centromeres in cen-Suv420-GFP expressing cells. **D)** Expression of cen-INCENP-mCherry does not limit cen-Suv420-GFP localization to centromeres. GFP-staining intensity between ACA peaks was measured for 3 kinetochore pairs in each of 30 metaphase cells per condition, in each of 3 biological replicates. **E)** expression of cen-INCENP-mCherry reduces the frequency of lagging chromosomes seen in cen-Suv420-GFP expressing anaphase cells. A minimum of 30 anaphase cells were scored per condition, for each of 3 biological replicates. **: p < 0.01, ***: p < 0.001. **F)** Quantification of area under the dose response curve (AUC) for each drug in cancer cell lines from the TCGA sorted by top and bottom quartile of either Suv39 or Suv420 expression shows that cells with high expression of either Suv39 or Suv420 exhibit increased sensitivity to Aurora kinase inhibition. Each dot represents an individual cancer cell line tested. Open circles indicate the mean of IC50 AUCs for each condition. *: p < 0.00625, **: p < 0.00125, ***: p <0.000125. Error bars are +/- SEM.

To explore this relationship more broadly, we utilized the drug sensitivity data described by the Welcome Sanger Institute (Yang et al., 2013) and RNAseq expression data from the Cancer Cell Line Encyclopedia that includes 1,457 cancer cell lines (Ghandi et al., 2019). Cell lines were sorted based on Suv39 or Suv420 expression level and then drug sensitivity of the top and bottom quartiles compared. We found that cell lines likely to have high H3K9 and/or H4K20 methylation states (due to high expression of Suv39 or Suv420) exhibit increased sensitivity to 5 out of 6 inhibitors that preferentially target Aurora kinase activity (Alisertib, CD532, GSK1070916, Genentech-CPD-10, and ZM447439) (Fig. 7F). This relationship does not reflect a general sensitivity to mitotic poisons as inhibition of the mitotic kinase MPS1 (MPS1IN1, TK3146) do not indicate a similar correlation with expression of these epigenetic regulators.

## Discussion

Here we demonstrate that Aurora B recruitment to centromeres is sensitive to changes in the epigenetic regulation of pericentromeric heterochromatin. We find that increased methylation of H3K9 and H4K20 corresponds with reduced Aurora B localization to centromeres and reduced phosphorylation of Aurora B substrates. Consistent with studies that directly perturb Aurora B function (Kallio et al., 2002, Hauf et al., 2003, Ditchfield et al., 2003), we show that such changes in centromere levels of the methyltransferases Suv39 or Suv420 are sufficient to stabilize kinetochore microtubule dynamics, limit merotelic error correction, increase lagging chromosomes during anaphase, and promote whole chromosome segregation errors.

### Suv39 and Suv420 overexpression contribute to increased aneuploidy

Analyses presented here and previously reported by The Meyerson group (Taylor et al., 2018) indicate a moderate but highly significant correlation between degree of aneuploidy and the independent expression levels of Suv39h1, Suv39h2, Suv420h1, and Suv420h2. Consistent with previous reports, our experimental data indicate that these isoforms of Suv39 and Suv420 also share at least partially overlapping roles in heterochromatin regulation and mitotic fidelity (Tsang et al., 2010, O’Carroll et al., 2000). As such, we expect that misexpression of any one of these enzymes (Suv39h1, Suv39h2, Suv420h1, or Suv420h2) could be sufficient to promote segregation errors and that the correlation of an individual enzyme with degree of aneuploidy would therefore be lower than if the function were served by a single enzyme. Though the correlation with aneuploidy for each enzyme is moderate, given the high significance of this relationship, we propose these correlations are consistent with a model whereby high expression levels of a Suv39 or Suv420 isoform are sufficient to promote mitotic defects and contribute to the generation of aneuploidy. Our data does not argue against the possibility that Suv39 or Suv420 function outside regulation of pericentromere heterochromatin may compromise genome stability in cancer. However, our experimental data showing that centromere enrichment of Suv420/H4K20me3 is sufficient to compromise mitotic fidelity, together with sustained pericentromere enrichment of H4K20me3 in cancer cells with high expression of Suv420, suggest that corruption of centromere regulation may be a contributing factor.

### Short term changes in centromere H3K9me3 and/or H4K20me3 corrupt mitotic fidelity without apparent defects in gross centromere structure

Work from other groups has described that a persistent increase in H3K9me3 over many cell cycles limits CENP-A deposition at the centromere on a human artificial chromosome (HAC), thus compromising centromere maintenance (Ohzeki et al., 2012, Martins et al., 2016, Martins et al., 2020). However, while induction of cen-Suv39-GFP of cen-Suv420-GFP for 24h (less than two cell cycles) is sufficient to cause dramatic changes in centromere levels of H3K9me3 and/or H4K20me3, it appears insufficient to compromise centromere formation or maintenance as we detect no change in overall levels of CENPA at the centromere, or in the outer kinetochore protein Hec1 (Supplemental Figure 5, Figure 6). Consistent with the sustained localization of critical centromere and outer kinetochore proteins, we find that cells expressing cen-Suv39-GFP or cen-Suv420-GFP remain competent to form stable end-on microtubule attachments that are sufficient to drive chromosome movement and segregation. However, we find that short-term increases in Suv39 and/or Suv420 at centromeres compromised CPC localization and reduced phosphorylation of Aurora B substrates, suggesting that spreading of the respective heterochromatin marks can impact centromere function in two distinct ways: first by subtly moderating regulation of microtubule attachments, and then more crudely by preventing centromere formation and maintenance. As methylation levels likely increase with duration of the experimental perturbation, these distinctions may arise due to experimental differences in the total amount of H3K9/H4K20 methylation achieved at the centromere, or instead reflect a cumulative impact of persistent epigenetic deregulation over several cell cycles remains unclear.

### Both direct and indirect increases in H4K20me3 levels corrupt mitotic fidelity

In otherwise normal cells, depletion of KDM4A, or overexpression of Suv420 alone promotes aberrant methylation throughout the genome; however changes in H4K20me3 at the centromere are moderate (Figure 3 B, (Manning et al., 2014)), suggesting that additional regulatory pathways may function to limit Suv420 enrichment or otherwise restrict H4K20me3 at centromeres. Nevertheless, this regulation of centromere/ pericentromere Suv420 localization may not be as tightly controlled in cancer contexts as we find that H4K20me3 remains enriched at centromeres in a panel of breast cancer cell lines that highly express Suv420 (Supplemental Figure 1). Furthermore, our study demonstrates that centromere targeting of Suv420 is sufficient to enhance H4K20me3 levels at centromeres without increasing H4K20me3 along chromosome arms. In this way, while we can not rule out that H3K9me3 independently impairs Aurora B localization at centromeres, our data suggest that increased Suv420 and/or H4K20me3, independent of Suv39/H3K9me3, is sufficient to restrict Aurora B localization and compromise mitotic fidelity. Importantly, expression of centromere-targeted GFP, even at levels higher than that of cen-Suv39-GFP or cen-Suv420-GFP, is insufficient to alter centromere H4K20me3 or Aurora B localization (Figures 3B & 4B). Together, these data suggest that Aurora B localization and mitotic fidelity are not generally perturbed by protein tethering to the centromere but are instead specifically sensitive to Suv420 and/or H4K20me3 levels.

Our data indicate that KDM4a depletion may disrupt Aurora B localization without a significant change in centromere H4K20me3 levels (Figure 3). Unlike expression of the cen-targeted constructs whose GFP tag allows us to confirm that each mitotic cell/centromere analyzed has overexpressed the enzyme, we do not have a similar report of KDM4A depletion at the single cell level and can not rule out the possibility that changes in methylation/Aurora B staining at centromeres may be impacted by variability of KDM4A depletion with the cell population. Nevertheless, our rescue experiments showing that inhibition of Suv420 methyltransferase activity (with A196) incompletely restores Aurora B localization (Figure 5) is consistent with Aurora B localization being at least partially independent of H4K20me3 levels.

HP1 directly binds the Aurora B-containing CPC through INCENP, and in doing so enhances the enzymatic activity of Aurora B (Abe et al., 2016, Kang et al., 2011). In turn, Aurora B-dependent phosphorylation of H3S10 limits HP1 association with H3K9 methylation (Fischle et al., 2005, Hirota et al., 2005), such that the interaction between HP1 and Aurora B both positively (through functional regulation) and negatively (through reduction of HP1 recruitment) regulates Aurora B activity at the centromere. Suv420 is similarly recruited to the pericentromere through interactions with HP1 and our data raise the possibility that CPC interaction with HP1 is limited when Suv420 is bound. Consistent with this possibility, we find that centromere-localization of the CPC is disrupted by centromere tethering of Suv420, but not overexpression of non-tethered Suv420. Interestingly, if true, this model would suggest that Suv420’s role in regulating CPC localization were in part independent of its methyltransferase activity.

### Disruption of heterochromatin boundaries impair centromere cohesion, transcription, and function

The constitutive heterochromatin marks H3K9me3 and H4K20me3 are enriched at pericentromeres but normally restricted from the core of the centromere (Sullivan and Karpen, 2004). This dynamic boundary between centromeric and pericentromeric heterochromatin is defined by H3K9 methylation and its disruption impacts transcription of the underlying DNA (Lam et al., 2006). In our experiments, the Suv39 and Suv420 fusion constructs target methyltransferase activity to centromeres by exploiting the CENP-B DNA binding domain. This binding domain recognizes a 17bp motif at centromeres to which the constitutive centromere protein CENP-B localizes (Muro et al., 1992). In recruiting Suv39 and Suv420 to this domain H3K9 and H4K20 methylation increases at ACA-stained centromeres indicating that heterochromatin, and potentially heterochromatin-associated proteins, has spread from the pericentromere into the centromere. Consistent with spreading of heterochromatin, we find levels of centromere and pericentromere transcripts are reduced following tethering of Suv39 or Suv420 to mitotic centromeres (Supplemental Figure 6C). In future studies, it will be important to determine if CPC recruitment and centromere function is generally sensitive to increased Suv420 at centromere/pericentromere regions, or specifically sensitive to the spreading of the repressive H4K20me3 mark into the centromere, where it is normally absent.

Early in mitosis Aurora B is localized along chromosome arms and must be re-localized to centromeres to ensure accurate chromosome segregation. Various regulatory mechanisms for recruitment and retainment of Aurora B to centromeres have been identified including positive regulation by phosphorylation of H3T3 and H2AT120 (Yamagishi et al., 2010, Kelly et al., 2010, Wang et al., 2010, Tsukahara et al., 2010). However, our data indicate that neither H3T3 phosphorylation nor H2AT120 phosphorylation are reduced when H4K20me3 is increased, making it unlikely that H3K9 or H4K20 methylation impinge on these regulatory mechanisms.

Instead, our data suggest that disruption of Aurora B localization following targeted recruitment of Suv39 or Suv420 to the centromere may arise due to misregulation of centromere cohesion and/or transcription. As with marks of heterochromatin, cohesin is enriched at pericentromeres (Eckert et al., 2007, Glynn et al., 2004, Weber et al., 2004). This distribution of cohesin is required for transcription of mitotic centromeres such that disruption of either pericentromere cohesin enrichment, or transcription itself, impairs Aurora B localization (Jambhekar et al., 2014, Kleyman et al., 2014, Perea-Resa et al., 2020). Work from our group and others have demonstrated that Suv420-dependent H4K20me3 enhances pericentromeric cohesin (Hahn et al., 2013, Manning et al., 2014, Bernard et al., 2001). This relationship would predict that Suv420-dependent methylation should promote centromere transcription and Aurora B localization. However, we find that following Suv39/Suv420 tethering to centromeres, increased centromere H4K20me3 corresponds with reduced, not enhanced, transcript abundance and Aurora B localization at centromeres (Supplemental Figure 6C). Together these data support a model whereby suppression of centromere transcription itself and/or the reduction in centromere transcripts resulting from spreading of H4K20me3 from the pericentromere into the centromere may be sufficient, irrespective of cohesin enrichment, to disrupt CPC localization.

### Increased H4K20me3 contributes to chromosome instability

The aneuploid chromosome content that results from mitotic segregation errors contributes to infertility, and is a primary cause of non-viable embryos and birth defects (Baker et al., 2009, Choi et al., 2009, Gao et al., 2007, Heilig et al., 2010, Kuukasjarvi et al., 1997, McClelland et al., 2009, Nowell, 1976, Rajagopalan and Lengauer, 2004, Swanton et al., 2009, Sotillo et al., 2010, Cucco and Musio, 2016). Persistent underlying defects in chromosome segregation, termed chromosome instability (CIN), (Baker et al., 2009, Sotillo et al., 2007, Weaver et al., 2007, Hagstrom and Meyer, 2003) are prevalent in cancer contexts where CIN promotes intratumor heterogeneity that in turn contributes to tumor evolution and drug resistance (Baker et al., 2009, Choi et al., 2009, Gao et al., 2007, Heilig et al., 2010, Kuukasjarvi et al., 1997, McClelland et al., 2009, Nowell, 1976, Rajagopalan and Lengauer, 2004, Swanton et al., 2009, Sotillo et al., 2010, Weaver et al., 2007). Merotelic kinetochore attachments are demonstrated to be the primary cause of segregation errors in human cells (Cimini et al., 2001, Cimini et al., 2003, Ganem et al., 2009, Thompson and Compton, 2008), suggesting that corruption of the Aurora B error correction pathway may be prevalent in CIN. Nevertheless, while mutations in genes directly involved in spindle structure, chromosome segregation, or mitotic checkpoints have been associated with a subset of cancers and hereditary disorders (Kim et al., 2012, Chung et al., 2012, Cahill et al., 1998, Cahill et al., 1999, Cucco and Musio, 2016), these mutations do not explain CIN in the vast majority of contexts (Wang et al., 2004, Negrini et al., 2010, Rajagopalan and Lengauer, 2004).

Our data suggest that changes in centromere methylation may underlie CIN in some cancer contexts. Consistent with this model, our work and others demonstrate that depletion of H3K9 demethylase enzymes, or high expression of H3K9 or H4K20 methyltransferase enzymes compromise mitotic fidelity in experimental systems and/or correspond both with aneuploidy and poor outcome in human cancer patients (Figure 1B, Supplemental Table 1, (Janssen et al., 2018, Black et al., 2012, Kupershmit et al., 2014, Frescas et al., 2008)). We find centromere tethering of either Suv39 or Suv420 is sufficient to compromise Aurora B kinase localization at centromeres and result in reduced phosphorylation of Aurora B substrates that are critical for proper chromosome segregation. Hec1, a key Aurora B substrate that governs kinetochore microtubule stability and mitotic error correction, is also a substrate for the related Aurora A kinase (DeLuca, 2017). Consistent with this redundant regulation, our data indicate that when centromere methylation is experimentally enhanced, or in cancer contexts where Suv39 or Suv420 expression is high and H4K20me3 is therefore likely high, sensitivity to inhibition of both Aurora A and Aurora B kinase is increased (Figure 7). The Aurora kinase family represent promising therapeutic targets that are actively being pursued in clinical and preclinical studies (Tang et al., 2017). Our results propose an intriguing possibility that levels of H4K20me3 (or expression of the enzymes that directly or indirectly promote this methylation state) may be predictive of CIN and may furthermore indicate sensitivity to molecular therapeutics that target Aurora B directly.

## Methods and Materials

### Cell Culture, siRNA and transgene expression

hTERT-RPE-1 (RPE-1) cells were grown in Dulbecco’s Modified Essential Medium (DMEM). Breast Cancer cell lines HCC1187, HCC202, and ZR-75-1 were grown in RPMI medium, SK-BR-3 were grown in McCoy’s 5a medium, and MCF7 grown in Eagles MEM medium. All cell culture medium were supplemented with 10% fetal bovine serum and 1% Penicillin + Streptomycin and cells grown at 37°C with 5% CO2.

Depletion of KDM4A was carried out through transient transfection of one of four individual, or a SMARTpool of all four, ON-TARGETplus siRNA constructs (horizon inspired cell solutions) using the RNAiMAX transfection reagent (Invitrogen) according to manufacturer’s instructions. Transfection with individual, or a SMARTpool of four, nontargeting siRNA sequences (horizon inspired cell solutions) were used as a negative control for KDM4A depletion. Knockdown efficiency was monitored by qPCR using KDM4A-specific primers. Expression of centromere-targeted, GFP-tagged cen-GFP, cen-Suv39-GFP and cen-Suv420-GFP fusion proteins, and non-targeted Suv420-GFP was achieved by cloning the respective cDNA into Addgene vector CENP-B DBD INCENP GFP (45237, Addgene) at NheI / BamH1. Inducible expression was achieved by cloning the GFP tagged constructs into plvx-Tre3G-IRES (631362, Clonetech) at Not1 / NdeI restriction cut sites. Expression vectors were transiently or stably expressed using Lipofectamine 3000 transfection reagent, according to manufacturer’s instructions. Inducible expression of transgenes was achieved by the addition of 2 μg/mL Doxycycline for 16-24h. “Mock” controls reflect the individual cell lines in the absence of Doxycycline-induction. Western blot analysis was used to confirm population level expression of each construct. Immunofluorescence for GFP was used to confirm expression and centromere localization for all single cell analyses performed. Sequences for all siRNA constructs and qPCR primers are represented in Supplemental Table 3.

### Metaphase spreads, fixation, and staining for immunofluorescence imaging

Metaphase spreads were prepared as in (Martins et al., 2016). Mitotic cells were collected by shake off following 3h incubation in 100ng/mL nocodazole, washed briefly in PBS (137mM NaCl, 2.7mM KCl, 10mM Na_2_HPO_4_, KH_2_PO_4_) and incubated in 75mM KCl for 12 minutes at 37°C. Cells were spun onto poly L lysine-coated chamber slides at 340xg for 5 minutes, then incubated in 37°C KCM buffer (120mM KCl, 20mM NaCl, and 10mM Tris HCl pH 8.0, in 0.1% Triton X-100) for 10 minutes.

For analysis of kinetochore protein localization, untreated (metaphase analysis) or nocodazole treated (prometaphase analysis) cells were prepared as in (Kleyman et al., 2014). Cells were incubated in 3.5% paraformaldehyde for 15min, quenched with 500mM ammonium chloride in PBS for 10min, incubated in ice cold methanol for 5min, and washed briefly with PBS. For cen-INCENP-mCherry expression, cells were transiently transfected with Addgene plasmid 45233 36h prior to doxycycline induction of cen-Suv420-GFP expression. For inhibition of Suv420 methyltransferase activity, cells were treated with 200nM A196 concurrent with doxycycline induction of cen-Suv420-GFP expression.

For metaphase and anaphase analyses of chromosome alignment and segregation, cells were fixed in 4% paraformaldehyde for 20min, washed briefly in PBS, post-extracted in PBS + 0.5% Triton X-100 for 10min, washed briefly in PBS, or alternatively fixed in ice cold methanol for 10min. For analysis of merotelic error correction capacity cells were incubated in the presence of 100 μM Monastrol for 4hrs, washed twice for 5min in drug-free growth medium at 37C, and incubated in growth medium supplemented with 25 μM MG132 (to prevent anaphase progression) prior to fixation at indicated times. For inhibitor sensitivity assays, cells were treated with indicated concentration of each drug for 12 hours prior to fixation.

Chromosome spreads were stained with antibodies diluted in KCM + 1%BSA. All other fixation methods were followed by 30min block in TBS-BSA (10mM Tris at pH 7.5, 150mM NaCl, 1% bovine serum albumin) + 0.1% Triton X-100 and incubation in primary then secondary antibodies in a humid chamber. Primary and secondary dilutions were made in TBS-BSA + 0.1% Triton X-100. DNA was detected with 0.2 μg/mL DAPI added to secondary antibody incubations. Coverslips were mounted onto slides using Prolong Antifade Gold (Molecular Probes).

### Antibodies

The following antibodies were used for immunofluorescence and/or western blot analyses: human anti centromere (ACA) (Antibodies Inc. 15-234); mouse anti H3K9me2 (ab1220, Abcam); rabbit anti H3K9me3 (ab176916, Abcam); rabbit anti H3 (ab1791, Abcam); rabbit anti H4K20me3 (ab9053, Abcam); rabbit anti H4 (ab7311, Abcam); rabbit anti H3S10p (04-817, Millipore); mouse anti α-tubulin (dm1α, Sigma); mouse anti Aurora B (AIM-1, BD Biosciences); rabbit anti H3T3p (97145, Cell Signaling); rabbit anti H2AT120p (61196, Active Motif); rabbit anti CENP-As7p (07-232, Upstate); rabbit anti CENP-A (2186, Cell Signaling); rabbit anti pHEC1S55p (pA5-85846, Invitrogen); mouse anti HEC1 (9G3.23, Novus Biologicals); goat anti GFP (ab6662, Abcam); rabbit anti GFP (2956, Cell Signaling); rabbit anti mCherry (PA5-34974, Invitrogen).

### Fluorescence Microscopy, Image Analysis, and Statistics

Cells were captured with a Zyla sCMOS camera mounted on a Nikon Ti-E microscope with a 60x Plan Apo oil immersion objective for fixed immunofluorescence, or a 20x CFI Plan Fluor objective for live cell imaging. All images within a single replicate were prepared and imaged in parallel, and images captured using the same exposure time. All single cells or single kinetochore measurements were normalized to the average intensity of the respective control condition and represented as fold-change in fluorescence. For metaphase and anaphase cell analyses, a minimum of 30 mitotic cells were analyzed per condition for each of 3 biological replicates. In these analyses, lagging chromosomes are defined as single chromosomes (marked by ACA-stained kinetochore) that are located between the two masses of segregating chromosomes (anaphase plates) and at least as far from the next nearest kinetochore as the anaphase plate is wide (based on ACA staining). Significance between biological replicates (n=3) was determined using a students’ two-tailed t-test. To assess centromere specific protein levels NIS Elements software was used to draw a line through ACA-stained centromere/kinetochore pairs. The intensity of antibody staining along the line was assessed for 3 kinetochore pairs per cell. To assess centromere enrichment, average pixel intensity of H4K20me3 staining was measured in 10 uniform ROIs defined at ACA-stained centromeres/kinetochores per cell and divided by the average H4K20me3 staining at ROIs defined on DAPI stained chromatin. Analyses were performed for a minimum of 30 cells in each of 3 biological replicates. Significance between biological replicates (n=3) was determined using a students’ two-tailed t-test. Live cell images were captured at 10 coordinates per condition at 5-minute intervals for 24hrs for each of 3 biological replicates. Mitotic timing was recorded from NEB until anaphase onset (Mercadante et al., 2019) for a minimum of 50 cells per replicate and significance between biological replicates (n=3) was determined using a students’ two-tailed t-test. All analyses were performed on unprocessed images. For publication images NIS elements deconvolution software was used. All images within a single panel were cropped comparably using ImageJ software to allow for direct comparison.

### qPCR analysis of centromere transcript abundance

RPE-1 cells carrying the inducible cen-Suv39-GFP or cen-Suv420-GFP construct were plated in cell culture flasks and induced, or not, with 2 μg/mL Doxycycline for 24h. 100 ng/mL Nocodazole was added to the growth medium for the last 4 hours of induction. Mitotic cells were collected via manual shake off. RNA was isolated with Trizol and purified using Qiagen’s RNAeasy columns. Purified RNA was quantitated by qPCR per the double delta Ct method that considers transcript changes relative to an internal control that is insensitive to the experimental manipulations (in this case GAPDH transcript), using transcript-specific primers (see Supplemental Table 3), a ABI 7500, and power up SYBR Green Master mix (Roche). Primers for centromere αsatellite transcripts were previously described (Liu et al., 2015, Xue et al., 2013).

### Gene Expression, Aneuploidy Correlation, Patient Survival, and Drug Sensitivity Analyses

For analyses of Suv39 and Suv420 isoform expression in cancer rna-seqv2 illumina hiseq RSEM normalized data was acquired through FireBrowse (firebrowse.org) and aneuploidy scores obtained from (Taylor et al., 2018). Both datasets were imported into an R statistical programming environment version 3.3.1 and run through a custom analysis pipeline to reformat and combine the datasets based on tumor sample. The Pearson correlation coefficient between aneuploidy score and gene expression was determined as described in (Taylor et al., 2018) and linear regression analysis performed to determine statistical significance where a p value of < 0.0025 reflects a significant correlation. RPKM-normalized expression values of Suv39 and Suv420 isoforms in normal and cancer contexts were obtained from FireBrowse and filtered to select and graph only cancer subtypes where paired normal tissue samples were available. The Kaplan-Meier Plotter platform (kmplot.com/analysis; (Nagy et al., 2018, Györffy et al., 2010)) was used to query survival data relative to Suv39 and Suv420 isoform expression to compare samples for top and bottom quartile expression for their respective genes. Hochberg’s step-up method corrected p values where determined and significance defined as p<= 0.0053. Gene expression data was downloaded from the Broad Cancer Cell Line Encyclopedia (CCLE) January 02, 2019 run containing expression data of 84,434 genes from 1,457 different cancer cell lines. Drug Screening IC50 AUC (area under the fitted dose response curve) data was downloaded from the Genomics of Drug Sensitivity in Cancer database GDSC1 run containing 518 different drugs across 988 different cell lines. Both sets of data were then imported into an R statistical programming environment version 3.3.1 and run through an analysis pipeline to reformat and combine the datasets based on cancer cell lines present in both. The data sets were sorted to independently identify the top and bottom quartile in terms of Suv39 or Suv420 expression. For both the high and low expression groups for each gene, the drug IC50 of each individual cell line and the mean +/- SEM of all cell lines in each group were graphed for each drug of interest. Significance was determined using a students’ two tailed t-test corrected for multiple comparisons using the Bonferroni correction method where p< 0.00625 is significant.

## Acknowledgements

We thank members of the Manning lab for critical reading and feedback on this manuscript. This work was supported by a Smith Family Award for Excellence in Biomedical Research and R00CA182731 to ALM.

The authors declare no competing interests.

## Author Contributions

AL Manning and CP Herlihy conceived and designed the experiments. CP Herlihy, S Hahn, E Crowley, and NM Hermance performed the experiments. AL Manning and CP Herlihy wrote the manuscript with input and approval from all authors.

