## Supplemental Figures and Legends for "Suv420 enrichment at the centromere limits Aurora B localization and function"

### Supplemental Figure Legends

**Figure S1. H4K20me3 is enriched near centromeres in breast cancer cells with high expression of Suv420.** Images and quantification of centromere localized H4K20me3 in a panel of breast cancer cell lines defined in the TCGA database as having high expression of Suv420. Kinetochore analyses were done on 3 kinetochores for each of 30 cells per condition. Scale bars are 5 μm.

**Figure S2. Depletion of KDM4A promotes H3K9 methylation and compromises mitotic fidelity. A)** Independent or pooled siRNAs sequences targeting KDM4A (see also Supplemental Table 3) result in similar level of mRNA depletion, as measured by qPCR. **B)** Quantification of the percent of control and KDM4A-depleted anaphase cells with one or more anaphase segregation defects for 30 anaphase cells for each of 3 biological replicates. **C)** Western Blot for GFP, Suv420, Suv39 following induction of expression of Suv420, cen-Suv420-GFP, and cen-Suv39-GFP constructs. Arrows indicate endogenous Suv39 and Suv420 protein. **D & E)** Representative images and quantification of single chromosomes from metaphase spreads 48 hours after KDM4A-depletion, mock or transient transfection of cen-Suv39-GFP or cen-Suv420-GFP show KDM4A depletion or cen-Suv39-GFP-expression, but not cen-Suv420-GFP expression, promotes an increase in centromere levels of H3K9me2 and H3K9me3 (left column of plots). Suv420-GFP does increase chromatin-associated levels of H3K9me3 (right column of plots; DAPI-localized staining, including centromeres). Quantification of H3K9me3 and H3K9me3 was performed across 10 paired ACA foci in each of 30 different metaphase spreads for each of 3 biological replicates. Total H3K9me3 staining was performed on DAPI-stained regions of each of 30 metaphase spreads for each of 3 biological replicates. Scale bar is 5 μm. For all panels, statistical analyses were performed between three biological replicates *: $p < 0.05$, **: $p < 0.01$. Error bars are +/- SD.

**Figure S3. Expression of cen-Suv39-GFP or cen-Suv420-GFP delays chromosome alignment. A)** Western blot analyses of GFP-tagged Suv420, CENP-B DNA binding domain alone (cen-GFP), cen-Suv420-GFP, and cen-Suv39-GFP showing that following induction, cen-GFP is expressed at levels higher than of the other fusion proteins. * indicates non-specific bands recognized by the GFP antibody. **B)** Representative images and quantification of metaphase alignment defects in cells induced to express cen-Suv39-GFP or cen-Suv420-GFP, compared to mock-induced cells. White arrowheads indicate single chromosomes that have failed to align. Error bars are +/- SD **C)** Quantification of Figure 3B showing cen-Suv420-GFP expression, but not Suv420-GFP promotes an increase in centromere enrichment of H4K20me3 (left column of plots), while only Suv420-GFP enhances total chromatin-localized H4K20me3 (right column of plots; DAPI-localized staining, including centromeres). Quantification of H4K20me3 was performed across 10 paired ACA foci in each of 30 different metaphase spreads for each of 3 biological replicates. Total H3K9me3 staining was performed on DAPI-stained regions of each of 30 metaphase spreads for each of 3 biological replicates. Scale bar is 5 $\mu$m. For all panels, statistical analyses were performed between three biological replicates *: $p < 0.05$, **: $p < 0.01$.

**Figure S4. Centromere localization of the CPC is reduced at centromeres of metaphase cells following depletion of KDM4A or expression of cen-Suv39-GFP or cen-Suv420-GFP.** Images and quantification of CPC component localization at centromeres of metaphase cells. Depletion of KDM4A or induced expression of centromere tethered cen-Suv39-GFP or cen-Suv420-GFP, but not untethered GFP-Suv420, reduced **A)** Aurora B and **B)** INCENP intensity at centromeres. Insets are 3X enlargements of single ACA-stained kinetochore pairs. Kinetochore analyses were done on 3 kinetochores for each of 30 cells per condition, per replicate **C)** qPCR analysis in cen-Suv39-GFP and cen-Suv420-GFP expressing mitotic cells show no change in transcript levels of CPC components Aurora B, Borealin, INCENP, and Survivin. For all panels statistical analyses were performed between three biological replicates, *: $p < 0.05$, **: $p < 0.01$. Error bars are +/- SD. Scale bars are 5 $\mu$m.

**Figure S5. Centromere tethering of Suv39 and Suv420 compromises phosphorylation of Aurora B substrates CENP-A and H3. A)** Images and quantification of metaphase cells showing that total centromere levels of CENP-A are not reduced following induction of cen-Suv39-GFP or induction of cen-Suv420-GFP. B) Images and quantification of metaphase cells showing that phosphorylation of

Aurora B substrate CENP-A (at serine 7) at the centromere is reduced following KDM4A depletion or induction of cen-Suv420-GFP expression. **C)** Western blot analysis of nocodazole-arrested mitotic cells induced to express cen-Suv39-GFP or cen-Suv420-GFP show no change in phosphorylation of Aurora B substrate H3 (at serine 10) Statistical analyses were performed between three biological replicates, *: $p < 0.05$. Scale bars are 5 $\mu$m.

**Figure S6. Increased H3K9 and/or H4K20 methylation reduce centromere transcription but does not compromise histone marks associated with mitotic Aurora B localization. A)** Images and quantification of centromere-levels of phosphorylated H3 (at threonine 3) and phosphorylated H2A (at threonine 120), epigenetic marks involved in localizing Aurora B to the centromere. Levels of H3T3p and H2AT120p are not altered following depletion of KDM4A or expression of cen-Suv39-GFP. **B)** Images and quantification showing that levels of H2AT120p are not altered following expression of cen-Suv39-GFP or cen-Suv420-GFP. Kinetochore analyses were done on 3 kinetochore pairs for each of 30 cells per condition, per replicate for each of 3 biological replicates. Scale bars are 5 $\mu$m. Error bars are +/- SD. **C)** qPCR analyses of repetitive centromere and pericentromere transcripts cen Sat-$\alpha$, D7Z1, D7Z2, and Sat-$\alpha$ from Chromosome 1 in nocodazole arrested mitotic cells show that centromere tethering of either cen-Suv39-GFP or cen-Suv420-GFP is sufficient to reduce transcript levels by more than half, while non-centromere transcripts (Supplemental Figure 3C) are unperturbed. Error bars are standard deviation between three biological replicates. Statistical analyses were determined by a student's t-test between replicates. *: $p < 0.05$

**Supplemental Table 1: Suv39 and Suv420 isoform expression exhibit a moderate but highly significant correlation with calculated aneuploidy score in several cancer subtypes.** Values reflect p values as determined by linear regression analysis. Shading indicates significance at $p < 0.0025$.

**Supplemental Table 2: Suv39 and Suv420 isoform expression in cancer inversely correlate with disease free survival.** The Kaplan-Meier Plotter platform (kmplot.com/analysis; (Nagy et al., 2018,

Györffy et al., 2010)) was used to query survival data relative to Suv39 and Suv420 isoform expression to compute samples for top and bottom quartile expression for their respective genes. Table reflects Hochberg's step-up method corrected p values for each relationship. Shading indicates significance of negative correlation at $p <= 0.0053$.

**Supplemental Table 3: Primers, Sequences, and Constructs used in this study**

Supplemental Figure 1

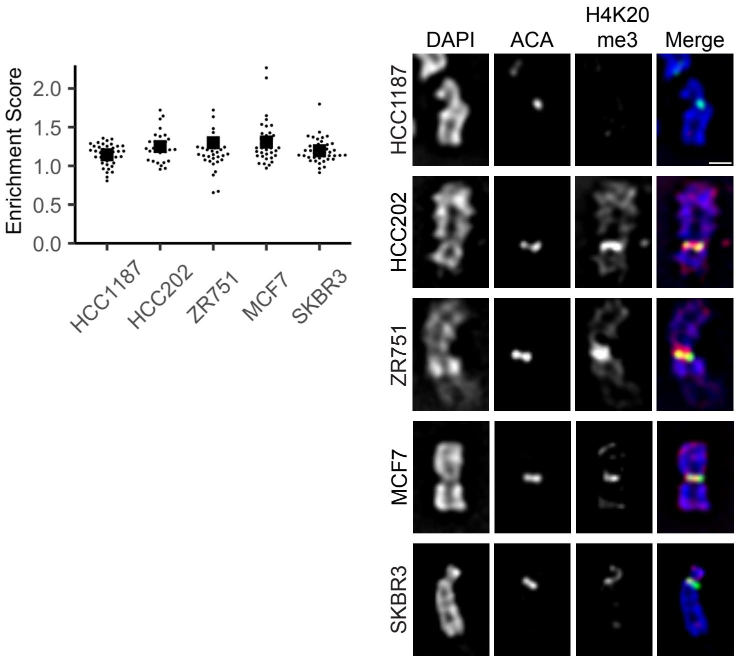

Supplemental Figure 2

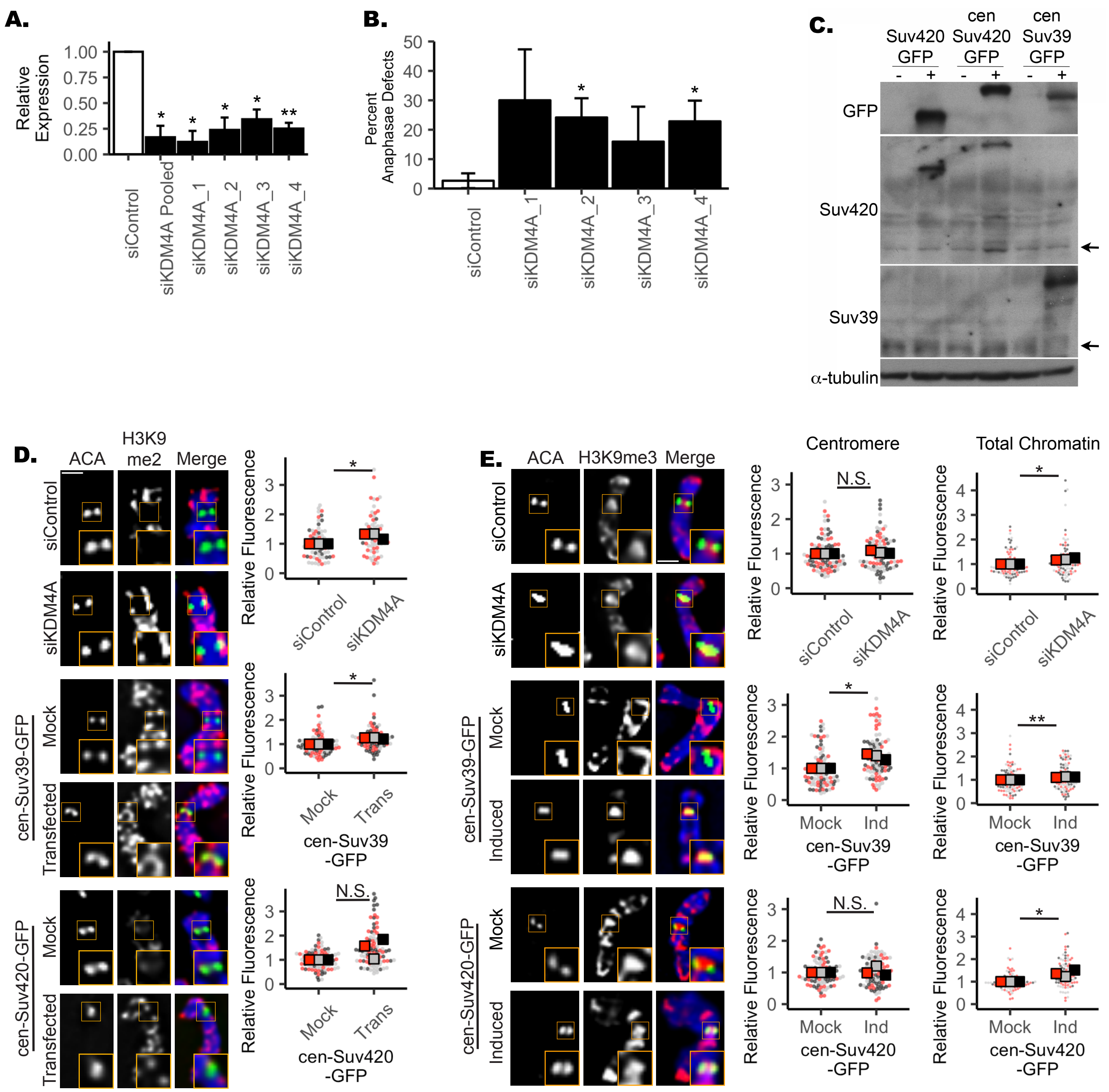

Supplemental Figure 3

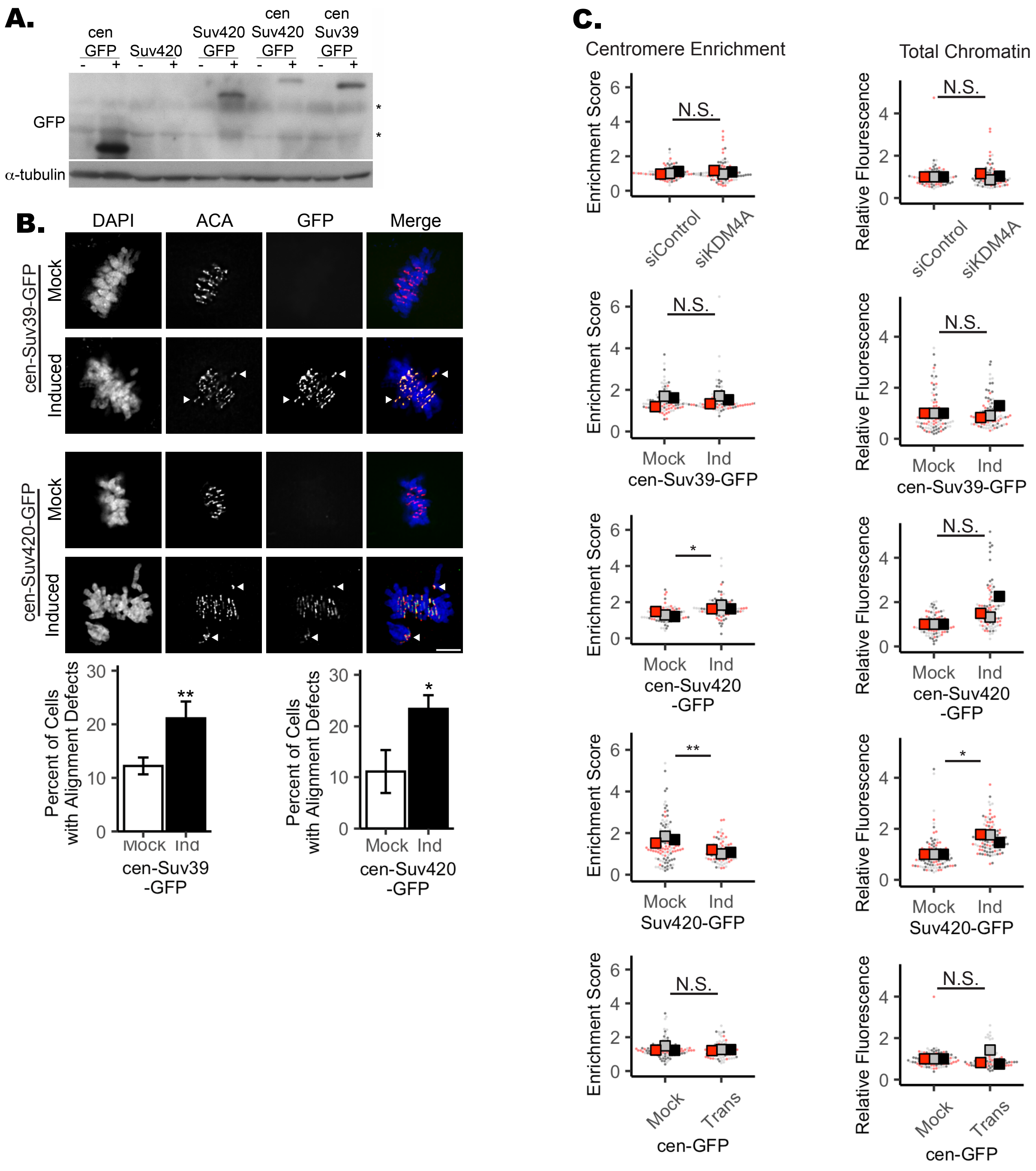

Supplemental Figure 4

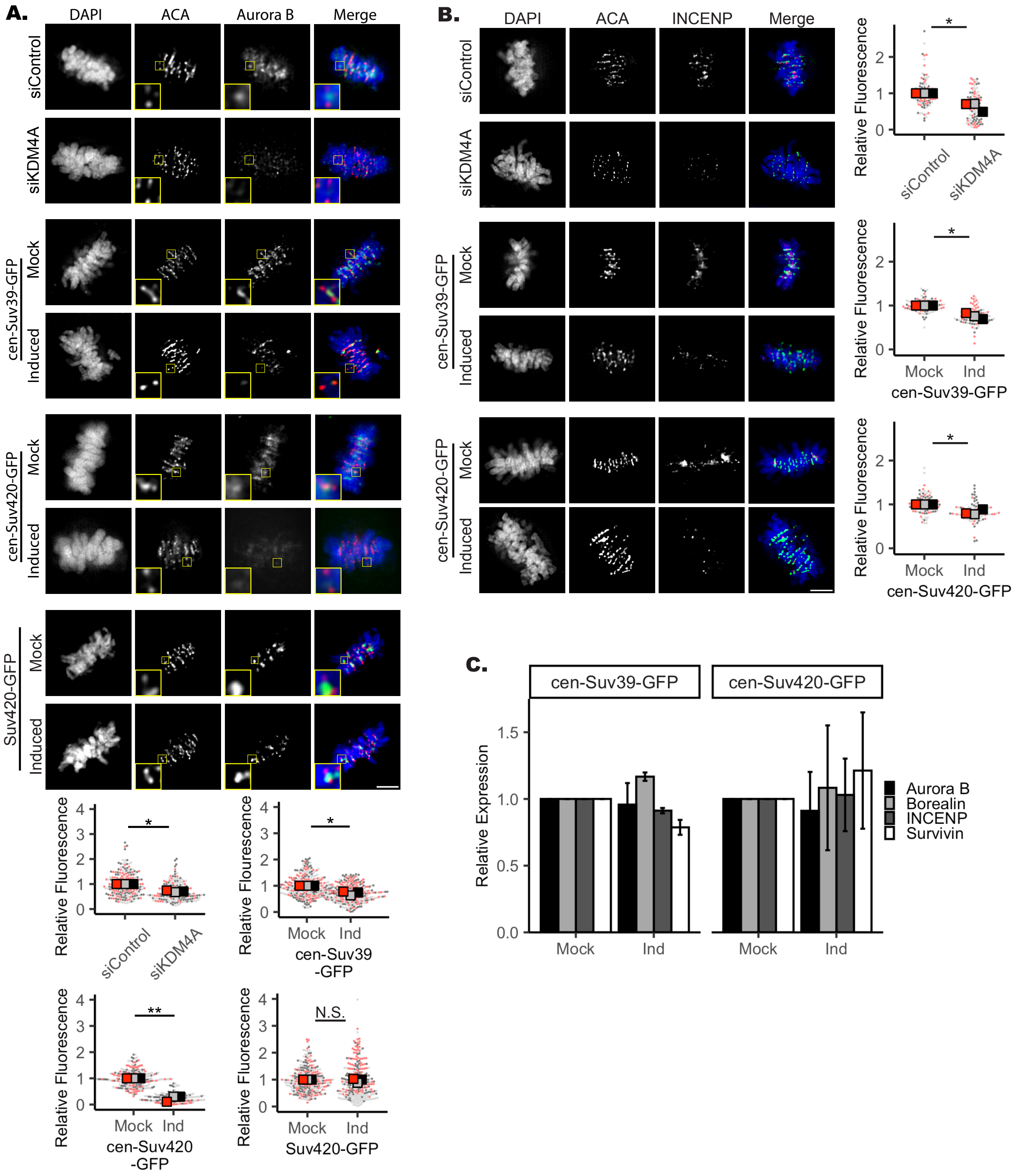

Supplemental Figure 5

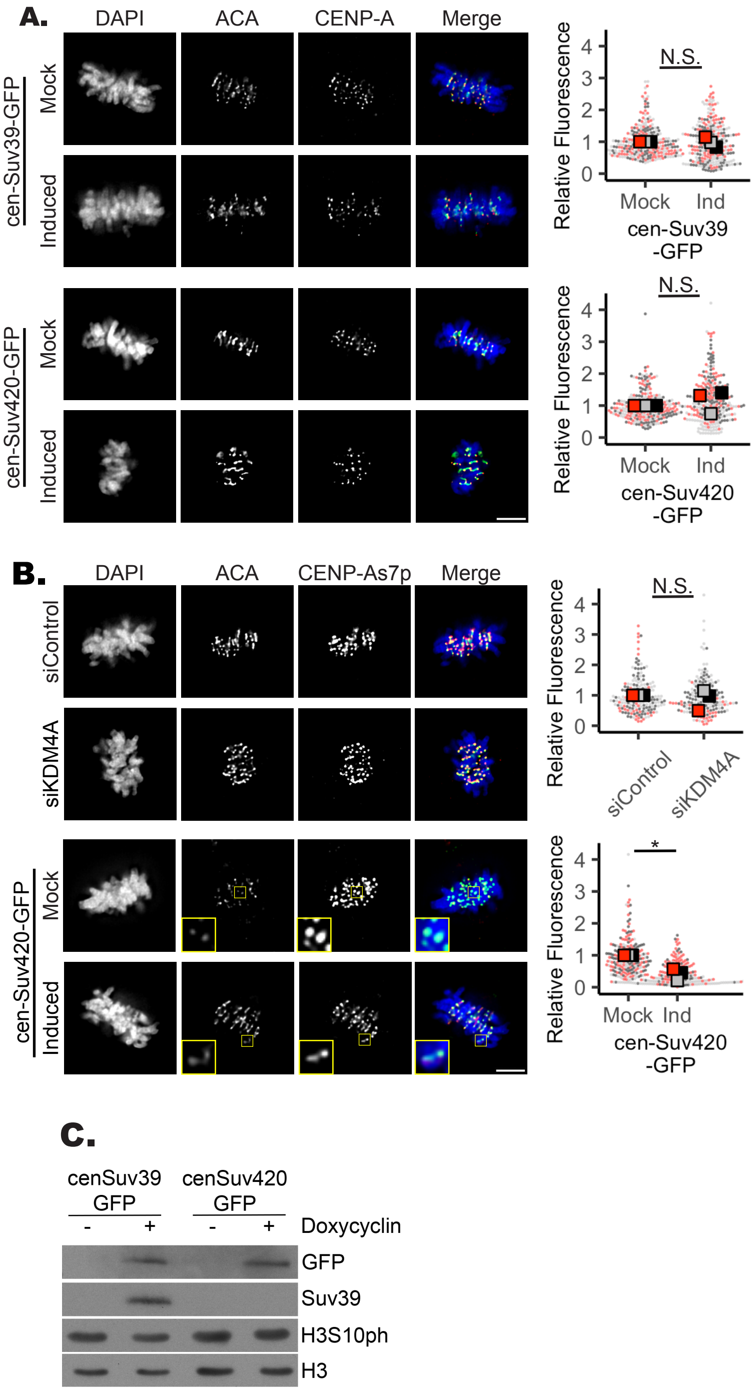

Supplemental Figure 6

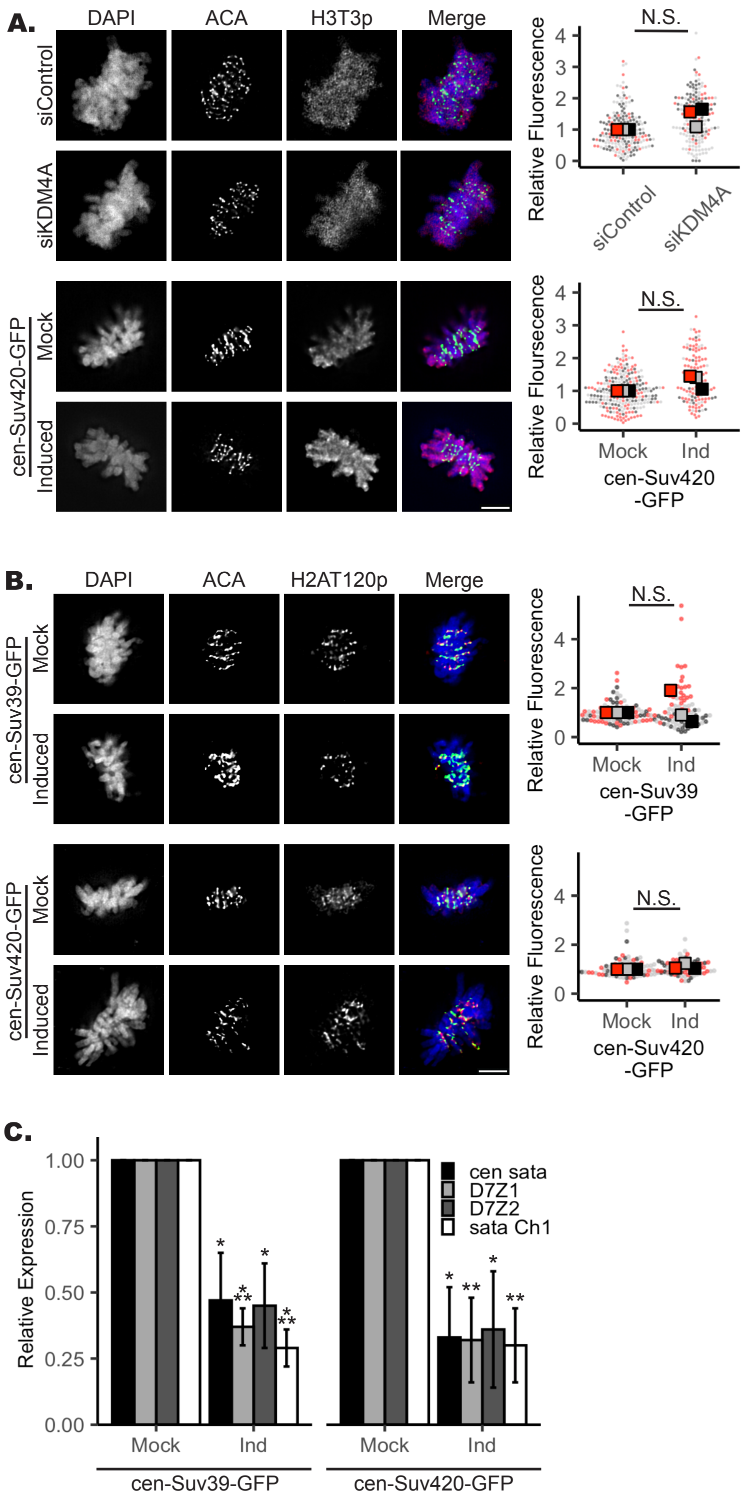

Supplemental Table 1

| Cancer Type | Abbreviation | Suv39h1 | Suv39h2 | Suv420h1 | Suv420h2 |
|---|---|---|---|---|---|
| Bladder Urothelial Carcinoma | BLCA | 0.0094 | 0.0000 | 0.2695 | 0.1560 |
| Breast Invasive Carcinoma | BRCA | 0.0000 | 0.0000 | 0.4494 | 0.3951 |
| Colon Adenocarcinoma | COAD | 0.0003 | 0.7974 | 0.3012 | 0.2998 |
| Esophageal Carcinoma | ESCA | 0.1506 | 0.1536 | 0.0622 | 0.0285 |
| Glioblastoma Multiforme | GBM | 0.0324 | 0.5125 | 0.0720 | 0.1038 |
| Kidney Chromophobe | KICH | 0.2305 | 0.8355 | 0.9010 | 0.0078 |
| Kidney Renal Clear Cell Carcinoma | KIRC | 0.0012 | 0.2833 | 0.0006 | 0.0008 |
| Kidney Renal Papillary Cell Carcinoma | KIRP | 0.1904 | 0.0603 | 0.8652 | 0.3673 |
| Glioma | LGG | 0.0000 | 0.0259 | 0.0000 | 0.0000 |
| Liver Hepatocellular Carcinoma | LIHC | 0.0000 | 0.0000 | 0.7800 | 0.0000 |
| Lung Adenocarcinoma | LUAD | 0.0000 | 0.0000 | 0.0001 | 0.0138 |
| Lung Squamous Cell Carcinoma | LUSC | 0.1520 | 0.0657 | 0.4101 | 0.5657 |
| Ovarian Serous Cystadenocarcinoma | OV | 0.0968 | 0.0655 | 0.0858 | 0.2250 |
| Prostate Adenocarcinoma | PRAD | 0.0000 | 0.3454 | 0.0240 | 0.0078 |
| Colorectal Adenocarcinoma | READ | 0.8655 | 0.0418 | 0.3215 | 0.2899 |
| Sarcoma | SARC | 0.0000 | 0.0010 | 0.0004 | 0.8933 |
| Skin Cutaneous Melanoma | SKCM | 0.2528 | 0.3194 | 0.8850 | 0.5028 |
| Stomach Adenocarcinoma | STAD | 0.0000 | 0.0000 | 0.0000 | 0.0000 |
| Thyroid Carcinoma | THCA | 0.0000 | 0.1053 | 0.2600 | 0.5170 |
| Uterine Corpus Endometrial Carcinoma | UCEC | 0.0000 | 0.0004 | 0.0002 | 0.0000 |

Significance cut-off at $p<0.0025$

Supplemental Table 2

| Cancer Type | Suv39h1 | Suv39h2 | Suv420h1 | Suv420h2 |
|---|---|---|---|---|
| Bladder carcinoma | 0.6133 | 0.3024 | 0.0206 | 0.5928 |
| Breast cancer | 0.8515 | 0.0173 | 0.4414 | 0.4606 |
| Cervical squamous cell carcinoma | 0.0092 | 0.512 | 0.0726 | 0.1118 |
| Esophageal adenocarcinoma | 0.4817 | 0.077 | 0.8126 | 0.0537 |
| Esophageal squamous cell carcinoma | 0.4504 | 0.0053 | 0.8196 | 0.3756 |
| Head-neck squamous cell carcinoma | 0.2241 | 0.3523 | 0.5863 | 0.1686 |
| Kidney renal clear cell carcinoma | 0.0188 | 0.8527 | 0.0002 | 1.70E-09 |
| Kidney renal papillary cell carcinoma | 0.2046 | 0.0011 | 0.4339 | 0.0841 |
| Liver hepatocellular carcinoma | 0.0533 | 3.10E-05 | 0.1269 | 0.2532 |
| Lung adenocarcinoma | 0.1725 | 0.2438 | 0.0631 | 0.5794 |
| Lung squamous cell carcinoma | 0.4053 | 0.3008 | 0.5597 | 0.2189 |
| Ovarian cancer | 0.4686 | 0.0641 | 0.9589 | 0.91 |
| Pancreatic ductal adenocarcinoma | 0.3382 | 0.5923 | 0.8592 | 0.0336 |
| Pheochromocytoma and Paraganglioma | 0.5412 | 0.0602 | 0.1547 | 0.7404 |
| Rectum adenocarcinoma | 0.3683 | 0.0973 | 0.5695 | 0.8821 |
| Sarcoma | 0.6517 | 0.0014 | 0.9985 | 0.1012 |
| Stomach adenocarcinoma | 0.4618 | 0.1844 | 0.8794 | 0.1555 |
| Testicular germ cell tumor | 0.1573 | 0.9879 | 0.9602 | 0.9307 |
| Thymoma | 0.0644 | 0.0045 | 0.079 | 0.0601 |
| Thyroid carcinoma | 0.7322 | 0.2861 | 0.8018 | 0.7307 |
| Uterine corpus endometrial carcinoma | 0.0262 | 0.026 | 0.1403 | 0.0097 |

Significance at p<= 0.0053

Supplemental Table 3

**Primers and siRNA Sequences**

| Primer Target | Forward Primer Sequence | Reverse Primer Sequence |
|---|---|---|
| GAPDH | CTAGCTGGCCCGATTTCTCC | GCGCCAATACGACCAAATCAGA |
| KDM4A | GGCTTTGGGCTGTAGATTCC | CCTAGCACTGGGATTCAGAGTT |
| Suv420h2 | CAACCATGACTGCAAACCCA | GCCGTAGAAGCATGTCACCT |
| Suv39h1 | CCTGCCCTCGGTATCTCTAAG | ATATCCACGCCATTTCACCAG |
| cen sat $\alpha$ | CATCACAAAGAAGTTTCTGAGAATGC TTC | TGCATTCAACTCACAGAGTTGAACC TTCC |
| D7Z1 | AAACGGGGTTTCTTCCTTTC | TGCCACAGCAAGAGTGTTTC |
| D7Z2 | AACCCCTTTGAGATGTGTGC | AGCTCCAAATGTCCAACTGC |
| Sat $\alpha$ Chr1 | TCATTCCCACAAACTGCGTTG | TCCAACGAAGGCCACAAGA |

**siRNA Sequences**

| Gene Target | Target Sequence | Source |
|---|---|---|
| Scr_1 | UGGUUUACAUGUCGACUAA | Dharmacon |
| Scr_2 | UGGUUUACAUGUUGUGUGA | Dharmacon |
| Scr_3 | UGGUUUACAUGUUUUCUGA | Dharmacon |
| Scr_4 | UGGUUUACAUGUUUUCCUA | Dharmacon |
| KDM4A_5 | GUAUGAUCUUCCAGACUUA | Dharmacon |
| KDM4A_6 | GCACGGACAUCAACCUUUC | Dharmacon |
| KDM4A_7 | GGGAUUCUAUCUCUUCUGA | Dharmacon |
| KDM4A_8 | GUGCGGAGUCUACCAAUUU | Dharmacon |

**Plasmids**

| Plasmid Name | Number | Source |
|---|---|---|
| CENP-B DBD INCENP GFP | 45237 | Addgene |
| CENP-B DBD INCENP mCherry | 45233 | Addgene |
| plvx-Tre3G-IRES | 631362 | Clontech |
| pLVX tet3G | 631191 | Clontech |
| cen-GFP | NA | This paper |
| cen-Suv39-GFP | NA | This paper |
| cen-Suv420-GFP | NA | This paper |
| Suv420-GFP | NA | This paper |
